# Structures of NF-κB p52 homodimer-DNA complexes rationalize binding mechanisms and transcription activation

**DOI:** 10.1101/2022.05.03.490500

**Authors:** Vladimir A. Meshcheryakov, Wenfei Pan, Tianjie Li, Yi Wang, Gourisankar Ghosh, Vivien Ya-Fan Wang

## Abstract

The mammalian NF-κB p52:p52 homodimer together with its cofactor Bcl3 activates transcription of κB sites with a central G/C base pair (bp), while it is inactive toward κB sites with a central A/T bp. To understand the molecular basis for this unique property of p52, we have determined its structure in complex with a P-selectin(PSel)-κB DNA (5’-GGGGT<u>G</u>ACCCC-3’) (central bp is underlined) and variants changing the central bp to A/T or swapping the flanking bp. The structures reveal a nearly two-fold widened minor groove in the central region of the DNA as compared to all other currently available NF-κB-DNA complex structures, which have a central A/T bp. Molecular dynamics (MD) simulations show free DNAs exist in distinct preferred conformations, and p52:p52 homodimer induces the least amount of conformational changes on the more transcriptionally active natural PSel-κB DNA in the bound form. Our binding assays further demonstrate that the fast kinetics driven by entropy is correlated with higher transcriptional activity. Overall, our studies have revealed a novel conformation for κB DNA in complex with NF-κB and suggest the importance of binding kinetics, dictated by free DNA conformational and dynamic states, in controlling transcriptional activation for NF-κB.

## Introduction

The binding of transcription factors (TFs) to their specific DNA response elements in the promoters/enhancers of target genes is the key event regulating gene transcription and consequent cellular processes. For proper gene expression, TFs must interact selectively at the correct place and time and assemble into high-order complexes with specific DNA sequences and cofactors (Natoli et al., 2005, Mulero et al., 2019). In eukaryotic genomes, the ability of TFs to select a small subset of relevant binding sites out of the large excess of potential binding sites within the genomes is the foundation upon which transcriptional regulation is built. Structural studies have provided valuable information on how various DNA binding domains recognize their cognate DNA binding sites at atomic resolution (Garvie and Wolberger, 2001). However, how TFs discriminate between closely related, but biologically distinct, DNA sequences is not well understood.

The NF-κB family of TFs regulates diverse biological responses (Zhang et al., 2017). Mammalian NF-κB is assembled combinatorially from five subunits, p50/NF-κB1, p52/NF-κB2, RelA/p65, c-Rel and RelB, into homo- and heterodimers which bind to specific DNA sequences, known as κB site or κB DNA. All five subunits share a highly conserved region at their N-termini, referred to as the Rel homology region (RHR), and the three-dimensional structures of the RHR are also highly conserved among these proteins. The RHR is roughly 300 residues in length and contains the N-terminal domain (NTD), dimerization domain (DD) and nuclear localization signal (NLS). The DD alone mediates protein homo- and heterodimerization; the NTD and DD together are responsible for DNA binding; the NLS region is flexible in solution and together with the DD forms the binding sites for the inhibitor of NF-κB (IκB) proteins.

The NF-κB proteins can be further divided into two sub-classes: the p50 and p52 subunits belong to class I by virtue of their lack of a transcriptional activation domain (TAD). The other three subunits, RelA, c-Rel and RelB, constitute class II with every member containing a TAD at its C-terminus. Mature p50 and p52 subunits are generated via incomplete proteolysis of their precursor proteins p105 and p100 (Supplemental Fig. S1A), respectively. Therefore, p50 and p52 possess a short glycine rich region (GRR) at their C-termini.

The initial discovery and characterization of several physiological κB DNAs established the pseudo-symmetric consensus sequence as 5’-(−5)G(−4)G(−3)G(−2)R(−1)N(0)W(+1)Y(+2)Y(+3)C(+4)C-3’ (Lenardo and Baltimore, 1989), where R = purines, N = any nucleotides, W = either A or T, and Y = pyrimidines. The subsequent identification of new NF-κB-DNA binding sites broadened the consensus to 5’-(−5)G(−4)G(−3)G(−2)N(−1)N(0)N(+1)N(+2)N(+3)C(+4)C-3’ (Chen and Ghosh, 1999, Mulero et al., 2019). The critical features of the consensus κB DNA sequence are the presence of a series of G and C bases at the 5’ and 3’ ends, respectively, while the bases at the central region can vary. X-ray structures of various NF-κB dimers in complex with different κB DNAs revealed conserved protein-DNA recognition modes for κB DNA that follows the consensus sequence (Muller et al., 1995, Ghosh et al., 1995, Cramer et al., 1997, Chen et al., 1998a, Huang et al., 2001, Moorthy et al., 2007, Fusco et al., 2009, Chen et al., 1998b, Chen et al., 2000, Escalante et al., 2002, Berkowitz et al., 2002, Chen-Park et al., 2002, Panne et al., 2007). The RHR of each monomer binds to half of a κB DNA, called half-site. A set of conserved amino acid (aa) residues mediate base-specific contacts to the 5’ and 3’ flanking G and C bases; the inner, more variable bases participate in important, but less base-specific interactions. The central bp lies at the pseudo-dyad axis of the dimer and is not directly contacted by the protein.

Genome-wide NF-κB-DNA motif identification studies revealed that NF-κB associates not only with consensus κB DNAs, but also with sequences containing only one half-site consensus, and even some sequences with no consensus (Lim et al., 2007, Martone et al., 2003, Zhao et al., 2014). *In vitro* binding experiments have been carried out to classify κB DNAs according to their binding specificity for different NF-κB dimers. The binding affinity displayed by various NF-κB dimers for distinct κB DNAs does not necessarily correlate with what occurs during regulation of gene expression *in vivo*. For example, the p50:RelA heterodimer binds tightly to most κB DNAs, whereas RelA:RelA and c-Rel:c-Rel homodimers bind many of the same sequences with relatively low affinity. However, detailed genetic experiments have shown that some genes are activated only in the presence of one or a subset of NF-κB subunits, such as mice lacking c-Rel exhibit defects in IL-2 and IL-12 expression (Kontgen et al., 1995, Hoffmann et al., 2003). In addition to specific gene activation, NF-κB dimers are also known to repress transcription. The RelA and p50 dimers have been shown to repress the expressions of *nrp1* gene, which is essential for osteoclast differentiation, and the interferon-stimulated response element (ISRE), respectively (Cheng et al., 2011, Hayashi et al., 2012). Both of these sites also display only half-site similarity to the κB DNA consensus. Structural and biochemical analyses of NF-κB-DNA binding have also revealed the existence of a large number of κB DNAs that display relatively similar affinities compared with κB consensus even though they lack one consensus half-site entirely (Ghosh et al., 2012, Siggers et al., 2011). Therefore, *in vitro* data do not fully capture the complexity of DNA recognition and gene regulation by NF-κB in cells.

NF-κB p52 is generated from the precursor protein p100 (Supplemental Fig. S1A), a tightly regulated process that requires specific stimuli. Unregulated p100 processing into p52 results in multiple myeloma and other lymphoid malignancies, which is detrimental to normal cellular function (Courtois and Gilmore, 2006, Annunziata et al., 2007, Keats et al., 2007). We previously demonstrated that the p52:p52 homodimer could sense a single bp change from G/C to A/T at the central position of a κB DNA (Wang et al., 2012). The p52:p52 homodimer binds both κB DNAs; but only in the case of the G/C-centric DNA, p52:p52 homodimer can associate with its specific cofactor Bcl3 (p52:p52:Bcl3 complex) and activate transcription by recruiting histone acetyltransferases. When bound to the A/T-centric DNA, the same p52:p52:Bcl3 complex represses gene transcription through the recruitment of histone deacetylases. It is intriguing that the identity of a non-contacted nucleotide should have such a drastic effect on transcriptional selectivity. Leung et al reported that the transcriptional activity of the RelA:RelA homodimer upon binding to A- and T-centric DNAs are different (Leung et al., 2004). Taken together, these reports strongly suggest that NF-κB transcriptional outcomes are coded within specific κB DNA sequences. Even small changes in the promoter specific κB DNAs, which do not alter the overall NF-κB binding affinity, might alter the gene expression profiles. Although structural studies have revealed a stereochemical mechanism of how NF-κB dimers bind κB DNAs, the effect of DNA conformation on complex formation remains underappreciated and it requires solid understanding of both structure and dynamics of κB DNAs to elucidate such a mechanism.

In the present study, we determined the crystal structures of the p52:p52 homodimer in complex with the PSel-κB DNA and two related DNAs where the central three positions were varied. PSel is a known target gene regulated by the p52:p52:Bcl3 complex in cells, and it contains a G/C-centric κB DNA in the promoter region (Pan and McEver, 1995, Wang et al., 2012). All three complexes revealed a widening of the DNA minor groove in the central region. *In vitro* experiments further demonstrated different thermodynamic and kinetic binding features: the binding of p52:p52 with transcriptional active promoters is driven by entropy with faster kinetics. The combination of structural, MD simulations, and biochemical studies presented here provides new insights into allosteric control by closely related κB DNAs on NF-κB-dependent transcriptional specificity.

## Results

### The central base pairs in PSel-κB DNA regulate p52:p52:Bcl3 transcriptional activity

Structures of several NF-κB dimers in complex with various κB DNAs have been reported over the past twenty-five years. In all these structures, the DNA sequences contain A/T-centric κB sites (Supplemental Fig. S2E) (Ghosh et al., 1995, Muller et al., 1995, Cramer et al., 1997, Chen et al., 1998b, Huang et al., 2001, Moorthy et al., 2007, Fusco et al., 2009, Chen et al., 1998a). The PSel-κB DNA (5’-GGGG<u>TGA</u>CCCC-3’) (the central bp is in red color, bps at ±1 positions are underlined), a natural binding site known to be specifically regulated by the p52:p52:Bcl3 complex (Pan and McEver, 1995, Wang et al., 2012), is distinctive from the canonical κB sites not only at the central position but also the two flanking positions. Whereas p50 and other subunits prefers an A:T at −1 and T:A at +1 positions, such as the MHC-κB site (5’-GGGG<u>ATT</u>CCCC-3’), PSel-κB contains T:A and A:T at the equivalent positions, respectively. We mutated the central and flanking bps to generate PSel (mutant A/T) and (−1/+1 swap) DNAs. Transcriptional activity of the p52:p52:Bcl3 complex was measured for these three and MHC-κB sites using a luciferase reporter based assay. The natural PSel luciferase reporter could be activated by endogenous NF-κB with co-expression of Bcl3 followed by LPS stimulation (Fig. 1A). To investigate the effects of PSel (mutant A/T) and (−1/+1 swap) on transcriptional activity, luciferase reporter constructs with the variants or MHC-κB site were co-transfected with p52 and Bcl3 expression plasmids. PSel (mutant A/T) showed 2-fold reduced reporter activity, while both PSel (−1/+1 swap) and MHC-κB showed drastically reduced transcriptional activity as compared to the natural PSel-κB (Fig. 1A; Supplemental Fig. S1C). These results suggest that the bp identity at all three positions in the central region are critical in determining transcriptional activity of the p52:p52:Bcl3 complex, which is in line with our previous study that the central bp of κB DNAs plays critical roles in transcriptional regulation (Wang et al., 2012).

**Figure 1.**
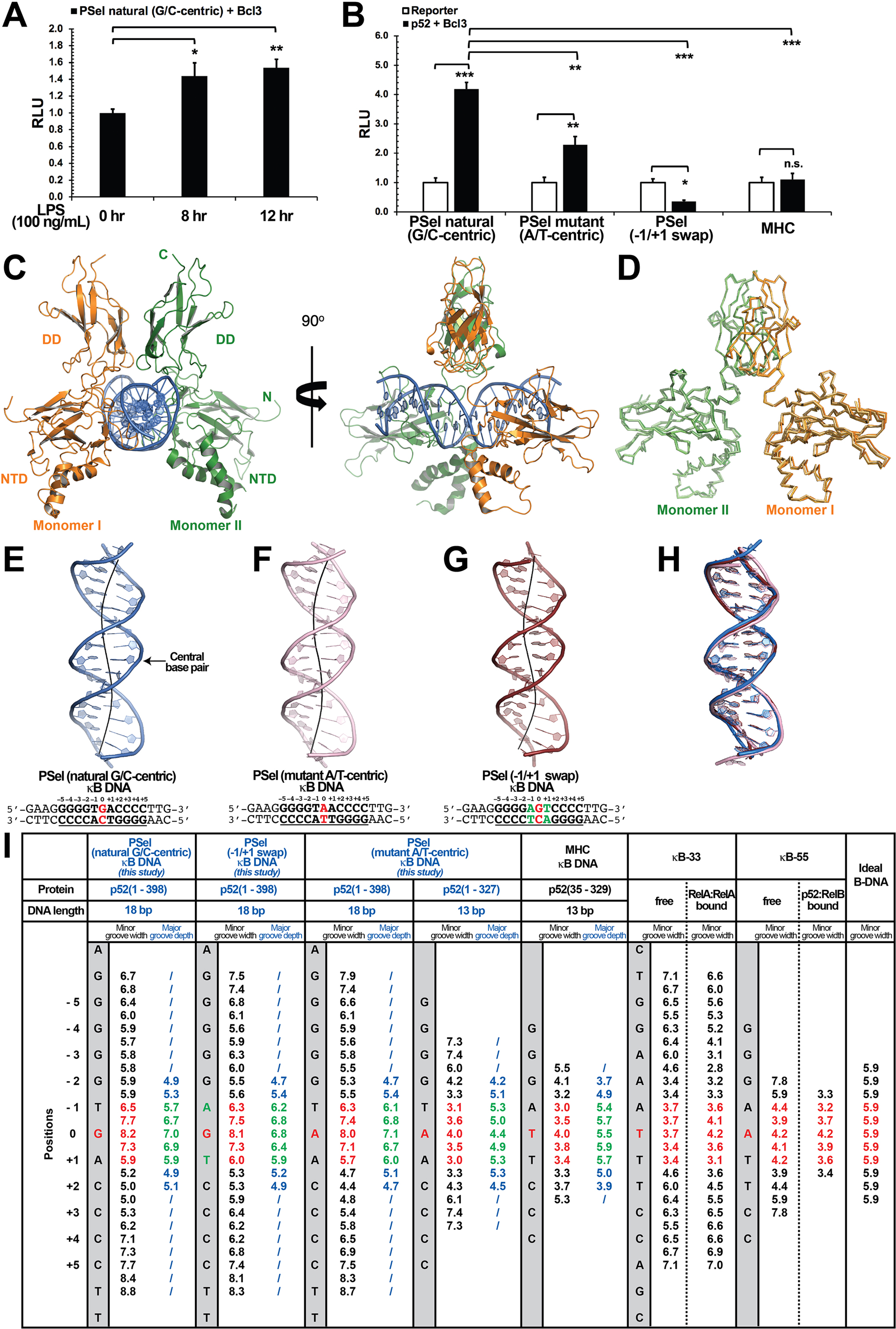
Crystal structures of p52:p52 homodimer in complex with PSel-κB DNA variants reveal distinct signatures. (A) The natural G/C-centric PSel luciferase reporter was activated by endogenous NF-κB with LPS stimulation and Bcl3 co-expression. The data were analyzed from three independent experiments performed in triplicate. RLU, relative luciferase unit. *p<0.05; **p<0.01 (t test). Error bars represent standard deviation (SD). (B) Luciferase reporter activity driven by co-expression of p52 and Bcl3 was reduced when the natural G/C-centric PSel site was mutated to A/T-centric or −1/+1 swap sites; and the MHC luciferase reporter was not activated by p52:p52:Bcl3 complex. The data were analyzed from three independent experiments performed in triplicate. *p<0.05; **p<0.01; ***p<0.001; n.s., not significant (t test). Error bars represent SD. (C) Overall structure of p52:p52 in complex with the natural G/C-centric PSel-κB DNA. (Left) Ribbon diagram showing the entire complex viewed down the DNA helical axis. The two p52 monomers are shown in orange (monomer I) and green (monomer II), respectively; and the DNA duplex is shown in blue; (Right) View of the complex after rotating 90° along the vertical axis. (D) Overlay p52:p52 homodimers in three PSel-κB DNA variants by their dimerization domain (DD). Monomer I is shown in tv_orange, bright orange and light orange; monomer II is shown in forest, tv_green and lime in the natural G/C-centric, mutant A/T-centric and −1/+1 swap complexes. All three structures are presented as backbone traces. (E-G) Structure of the 18bp PSel-κB DNAs with (E) natural G/C-centric (blue), (F) mutant A/T-centric (light pink) and (G) −1/+1 swap (ruby). The DNA bps as observed in the co-crystal structures are shown in filled sticks. The view is onto the central minor groove. The nucleotide sequences used in co-crystallization are shown at the bottom, with κB DNA underlined and numbering scheme indicated above; the central position 0 is highlighted in red, and the swap of −1 and +1 positions is highlighted in green. (H) Overlay of natural G/C-centric, mutant A/T-centric, and −1/+1 swap PSel-κB DNAs in (E-G). (I) Table showing minor groove widths and major groove depths (Å); the ideal B-form DNA was built using Coot program (Emsley and Cowtan, 2004, Emsley et al., 2010) based on the sequence of PSel-κB DNA. The minor groove widths at the central region from position −1 to +1 are shown in red, and the corresponding major groove depths are shown in green. Geometrical parameters and the helical axes were calculated with Curves*+* (Blanchet et al., 2011, Lavery et al., 2009).

### Widened minor groove in PSel-κB DNA in complex with NF-κB p52:p52 homodimer

Since only p52 mediates DNA interactions in the p52:p52:Bcl3 complex (Supplemental Fig. S1D) (Bours et al., 1993), we focused our study on (p52:p52)-DNA and speculated that the observed transcriptional differences could be due to different structural features of (p52:p52)-DNA complexes. We solved the crystal structures of p52:p52 homodimer in complex with all three PSel-κB DNAs (Fig. 1C-H; Table 1). The p52 protein works as a bridging factor between target DNAs and Bcl3; therefore, a recombinant p52 protein (aa 1-398) which could form complex with Bcl3 was co-crystallized with the DNAs (Supplemental Fig. S1E-H). This p52 construct contains most of the GRR region which was not included in any previous NF-κB structures (Supplemental Fig. S1B, S2E); however, no electron density was observed for the C-terminal part (aa 330-398) in the structures.

The overall structures of p52:p52 in complex with the natural PSel-κB DNA and two variants are similar to each other (Fig. 1D, H). However, compared to previously known structures of NF-κB-DNA complexes, two striking differences are observed. One is that all three PSel-κB DNAs exhibited a distinct widening of the minor groove at the two base-steps around the central position 0 (−1 to 0 and 0 to +1), with width of ∼7.5 Å (Fig. 1E-G, I). In comparison, the A/T-centric κB DNAs studied earlier, κB-33 (5’-GGAA<u>ATT</u>TCC-3’) (Chen et al., 1998b, Huang et al., 2005) and another one that we now name κB-55 (5’-GGG<u>AAT</u>TCCC-3’) (Moorthy et al., 2007, Fusco et al., 2009), have significantly compressed minor groove in both their bound and free states as compared to an ideal B-form DNA (Fig. 1I; Supplemental Fig. S2A-B). Compressed minor groove width (MGW) is a common feature of all A/T-centric κB DNAs bound to NF-κB dimers which is remarkably different from the MGW of the PSel-κB DNAs seen in the present structures (Supplemental Fig. S2E).

### The widened minor groove is observed with long p52 proteins

The other difference observed for the three p52:p52 structures reported here concerns the organization of the dimer and the complex with DNA. The p52-MHC-κB DNA (which is A/T-centric) complex is the only previously determined crystal structure of NF-κB p52:p52 homodimer (Cramer et al., 1997). Superposition of the p52:p52 homodimer in the MHC-κB and natural PSel-κB complexes aligned by the DDs reveals large rigid body movement of NTDs with rotation of ∼20° and translation along rotation axis of ∼1.4 Å (Fig. 2A). This results in shifting of the NTD along PSel DNA toward its flanks by ∼13 Å for both sides. In addition, the minor groove of the MHC-κB DNA at the central segment is compressed like all other NF-κB-DNA complexes indicated above (Fig. 1I; Supplemental Fig. S2C).

**Figure 2.**
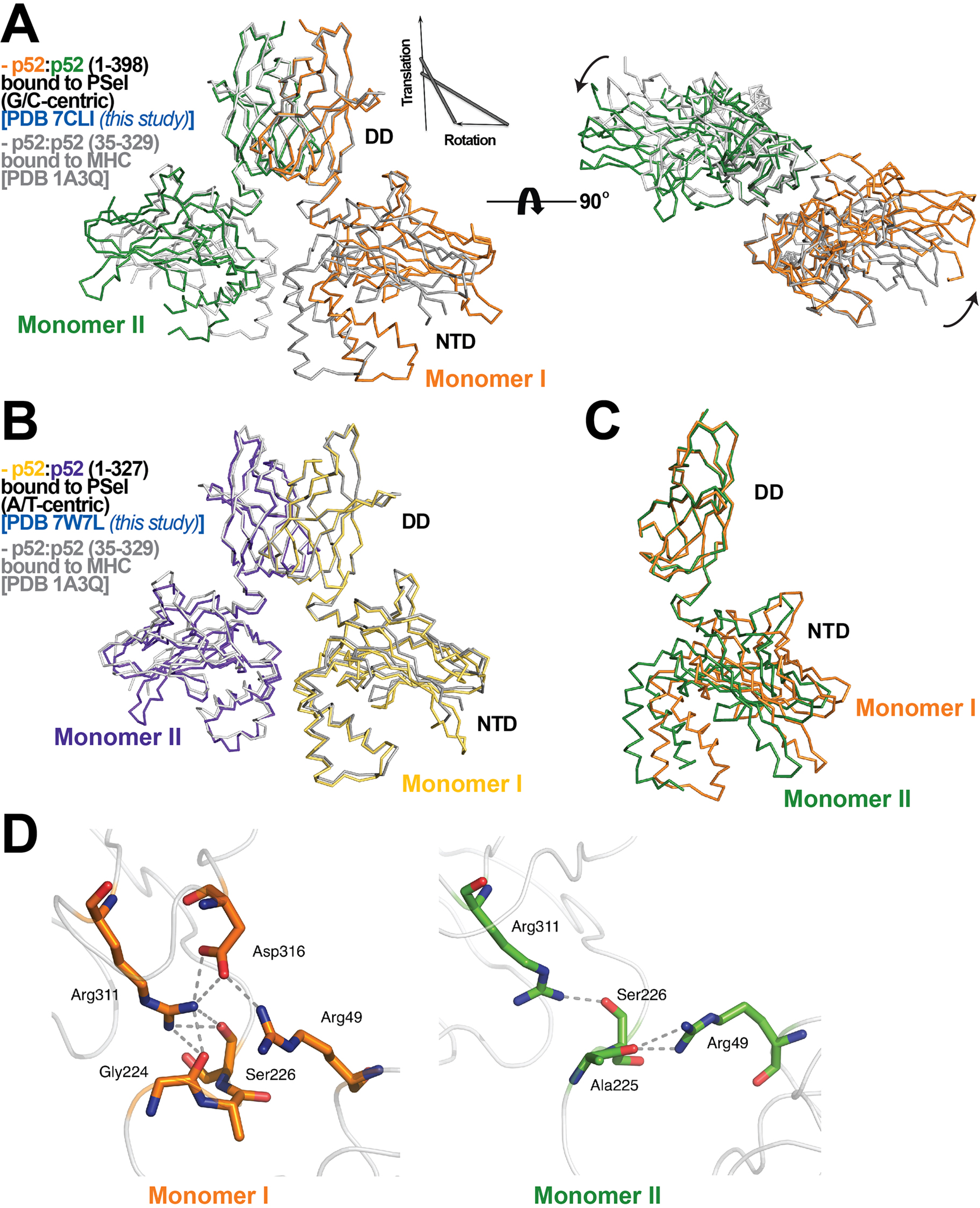
p52:p52 dimer conformations. (A) (Left) Overlay of p52:p52 (aa 1-398) in complex with the 18bp natural G/C-centric PSel-κB DNA (PDB 7CLI, this study)(in orange and green for monomer I and II, respectively) and p52:p52 (aa 35-329) in complex with the MHC-κB DNA (PDB 1A3Q)(in gray). Diagram explains rigid-body movement of the NTD. (Right) View of the complex after rotating 90° along the horizontal axis. Both protein structures are presented as backbone traces. (B) Overlay of the short p52:p52 (aa 1-327) in complex with the 13bp PSel(mutant A/T)-κB DNA (PDB 7W7L, this study)(in yelloworange and purpleblue for monomer I and II, respectively) and p52:p52 (aa 35-329) in complex with the MHC-κB DNA (PDB 1A3Q)(in gray). (C) Conformational differences between two p52 monomers in complex with the 18bp natural G/C-centric PSel-κB DNA (PDB 7CLI, this study) are shown by superposing their DDs. The two monomers are presented as backbone traces. (D) Hydrogen bonding network at the interdomain interface between DD and NTD in each p52 monomer. In monomer I, the two domains form contacts with each other through a wide network of H-bonds between the side chains of Arg49 from the NTD and Gly224, Ser226, Arg311 and Asp316 from the DD; whereases in monomer II, there are only contacts between Arg49 and Ala225, as well as Ser226 and Arg311.

The PSel-κB DNAs (18 bp) and recombinant p52 protein (aa 1-398, including the GRR) used in the current study are both longer than those in the MHC-κB DNA complex (13bp and aa 35-329). In fact, all currently available structures of NF-κB-κB DNA complexes in the Protein Data Bank (PDB) contain short NF-κB proteins (only NTD and DD) and A/T-centric κB DNAs (Supplemental Fig. S2E) (Ghosh et al., 1995, Muller et al., 1995, Cramer et al., 1997, Chen et al., 1998b, Huang et al., 2001, Moorthy et al., 2007, Fusco et al., 2009, Chen et al., 1998a). To test if the DNA and protein length variations induce structural changes in the complex, we crystallized a 13bp PSel (mutant A/T) DNA bound to a shorter p52 protein (aa 1-327). The conformation of this complex is nearly identical to p52-MHC-κB complex with MGW less than 4 Å at the central position (Fig. 2B, 1I; Supplemental Fig. S2D). This crystal with short p52 is in a different crystal form compared to the three structures with the long p52, and it is also in a different crystal form compared to the MHC-κB DNA complex, suggesting that crystal packing is unlikely to be the main cause of the structural differences, and that both the DNA and protein lengths play significant roles.

Therefore, our structures demonstrate a correlation between the length of the p52 protein and the conformation of the κB DNA and the organization of the p52:p52 dimer in the complex. As discussed above, the short p52 protein (aa 1-327) failed to interact with Bcl3 (Supplemental Fig. S1E-H), partly due to the lack of the GRR. We used the long p52 protein (aa 1-398) for the rest of the studies.

### Distinct protein-DNA interactions in the p52:p52-DNA complexes

The widening of the minor groove propagates from the central position to all four base steps on both sides with values around 5-6 Å (Fig. 1I). This widening and the consequent deepening of the major groove have significant impact on protein-DNA interactions. The most significant of which is the loss of cross-strand base contacts by Arg52 (Fig. 3A). The cross-strand contacts between the homologous Arg (Arg54 in p50 and Arg33 in RelA) and DNA is observed in all other A/T-centric NF-κB-DNA structures (Fig. 3B; Supplemental Table S1).

**Figure 3.**
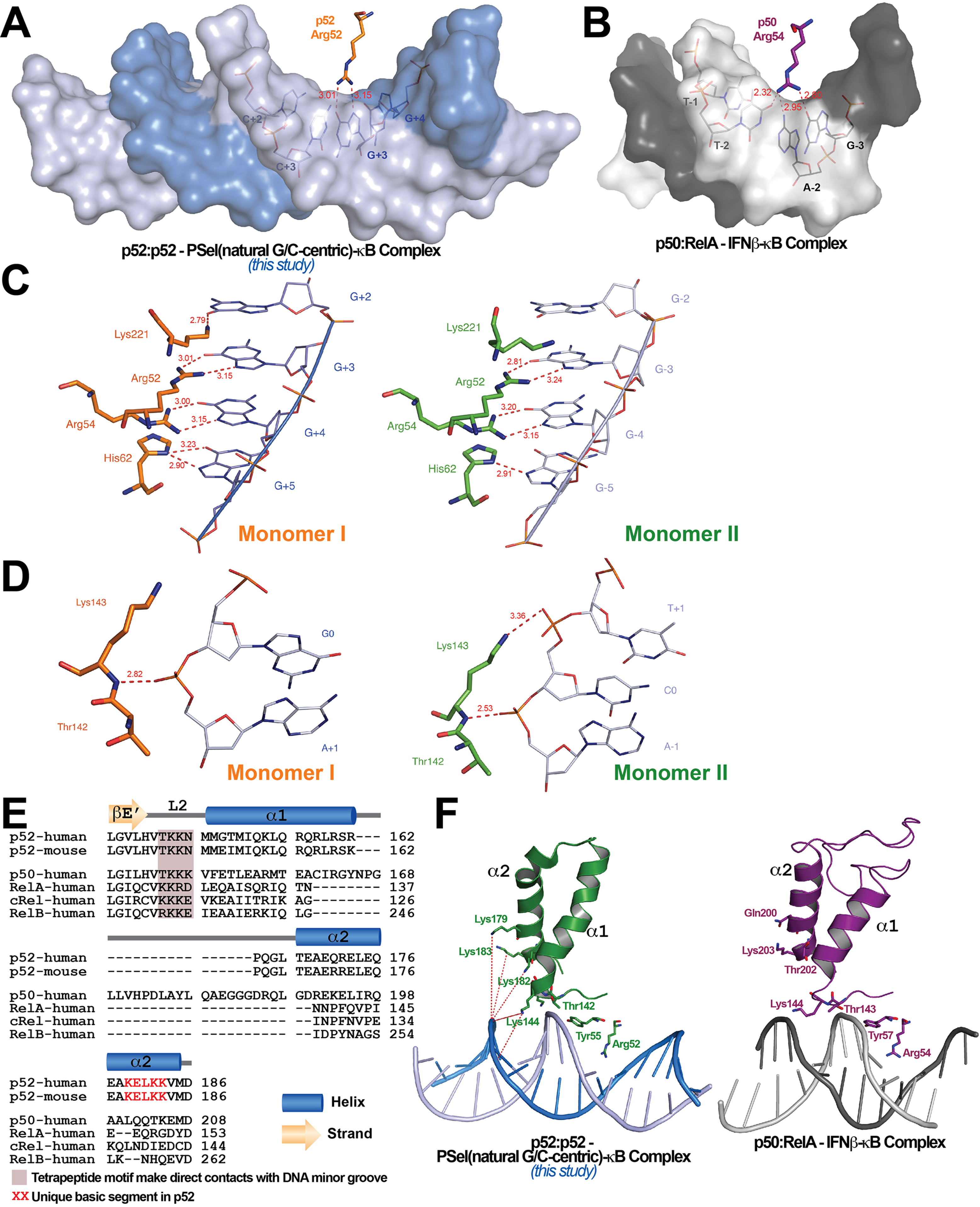
Protein-DNA contacts. (A) Arg52 of p52 in the PSel-κB complex (PDB 7CLI, this study) only makes base-specific contacts with G at +3 position. (B) The corresponding Arg54 of p50 in the p50:RelA-IFNb-κB complex (PDB 1LE5) makes base-specific contacts with A at −2 and G at −3 positions as well as additional cross-strand contacts with T at −2 position. (C) DNA based-specific contacts made by Arg52, Arg54, His62 and Lys221 of p52 (Left) monomer I and (Right) monomer II in complex with the natural PSel-κB DNA. H-bonds are indicated as red dotted lines with distances labelled. Noted that Lys221 in monomer II is in a different conformation and has no specific contacts with DNA. (D) DNA backbone contacts made by Lys143 of p52 (Left) monomer I and (Right) monomer II. (E) Sequence alignment showing the unique basic segment in p52 among all NF-κB family members. Both human and mouse sequences of the p52 subunit are shown. Only human sequences are shown for the rest of the family members. Secondary structures and connecting loops are drawn above the sequences. (F) (Left) The unique basic segment in p52 NTD helix a2 interacts with PSel-κB DNA in the present structure (PDB 7CLI, this study); (Right) these interactions are absent in p50 subunit in the p50:RelA-IFNb-κB complex (PDB 1LE5).

The other notable feature of the PSel-κB DNA complexes is the highly asymmetric DNA contacts by p52:p52 homodimer. Monomer I is closer to its cognate half-site making more direct contacts with the DNA than monomer II (Supplemental Fig. S3A). Although asymmetric DNA binding by the symmetric homodimer is a common feature in all NF-κB-DNA complexes, it is significantly more pronounced in the present structures. Moreover, the p52:52 homodimer also displays substantial asymmetry. With the DD of the two monomers in superposition, the NTDs rotate from each other by ∼6° (Fig. 2C). The interdomain interaction is extensive in monomer I compared to that in monomer II (Fig. 2D).

In the PSel-κB complex, the side chains of Lys221, Arg52, Arg54 and His62 in p52 monomer I make direct base-specific contacts to four consecutive G(s) from +2 to +5 positions (Fig. 3C, Left). In addition, Ser61 also makes direct contact with A at ±6 and ±7 positions; these contacts are not possible for the short p52 (aa 1-327) co-crystallized with 13bp κB DNAs such as MHC and PSel (mutant A/T)-κB (Supplemental Fig. S3A-B). p52 monomer II makes contact with only three G(s) from position −3 to −5 (Fig. 3C, Right). The conformation of loop L3 in the two p52 monomers are different; consequently, only Lys221 in monomer I makes specific contacts with G at position +2 (Supplemental Fig. S3C). Glu58 helps to position Arg52, Arg54 and His62, and makes base-specific interaction to the opposite C at ±3 position.

In addition to base-specific interactions, there are multiple protein contacts to the DNA phosphate backbone, mostly to the central region of the DNA. The side chain of Cys57 hydrogen bonds to the backbone phosphate group of C at ±2; and the side chains of Tyr55 and Lys143 hydrogen bond to the backbone phosphate group of A at ±1. Only in monomer II does the side chains of Lys143 make an additional hydrogen bond (H-bond) to the backbone phosphate group of C at position 0 (Fig. 3D). Interestingly, all other NF-κB-DNA complexes, including the short p52:p52 homodimer bound to both 13bp MHC-κB and PSel (mutant A/T)-κB DNAs, exhibit more backbone contacts by Gln284 and Gln254 (Supplemental Fig. S3B; Supplemental Table S1). The presence of an additional positively charged residue in loop L2 in the other NF-κB subunits (p52: T^142^KKN; p50: TKKK; and RelA: KKRD) enhances backbone binding at the minor groove side including cross-strand interactions (Fig. 3E). In addition, there is a unique basic segment in p52, a peptide rich in basic residues (K^179^ELKK), located near the end of helix α2 (Fig. 3E). These basic residues possibly mediate long-range electrostatic interactions with the negatively charged DNA backbone which might pull the DNA strands away from each other towards the p52 protein (Fig. 3F). In summary, amino acid composition in loop L2 and helix α2 might play an important role in determining DNA binding by the NF-κB dimers.

### MD simulations reveal free DNAs exist in distinct preferred conformations

In order to investigate whether MGW of the PSel-κB DNA variants observed in the current complexes is induced by the protein or it is intrinsic to DNA sequences, we carried out microsecond-long MD simulations for the four κB DNAs in free form. The simulations started from the DNA conformation in the crystal structures of the complexes, with the three PSel-κB DNA variants having a widened minor groove and the MHC-κB DNA having a narrow minor groove. At the end of the simulation, PSel (natural G/C-centric) and (mutant A/T-centric) maintained the widened minor groove, while PSel (−1/+1 swap) displays a narrow minor groove at the central 0 position similar to MHC-κB DNA in the simulations (Fig. 4A). The swap of T and A at ±1 positions reverses the geometric conformation of bps at both positions (Fig. 4C). Specifically, these swaps cause an opposite orientation of each nucleotide, forcing the bps to adopt an opposite shear and buckle direction compared to those on the non-swapped DNAs. The thymine at both positions slides and tilts towards the minor groove simultaneously, narrowing the central minor grooves (Supplemental Fig. S4A).

**Figure 4.**
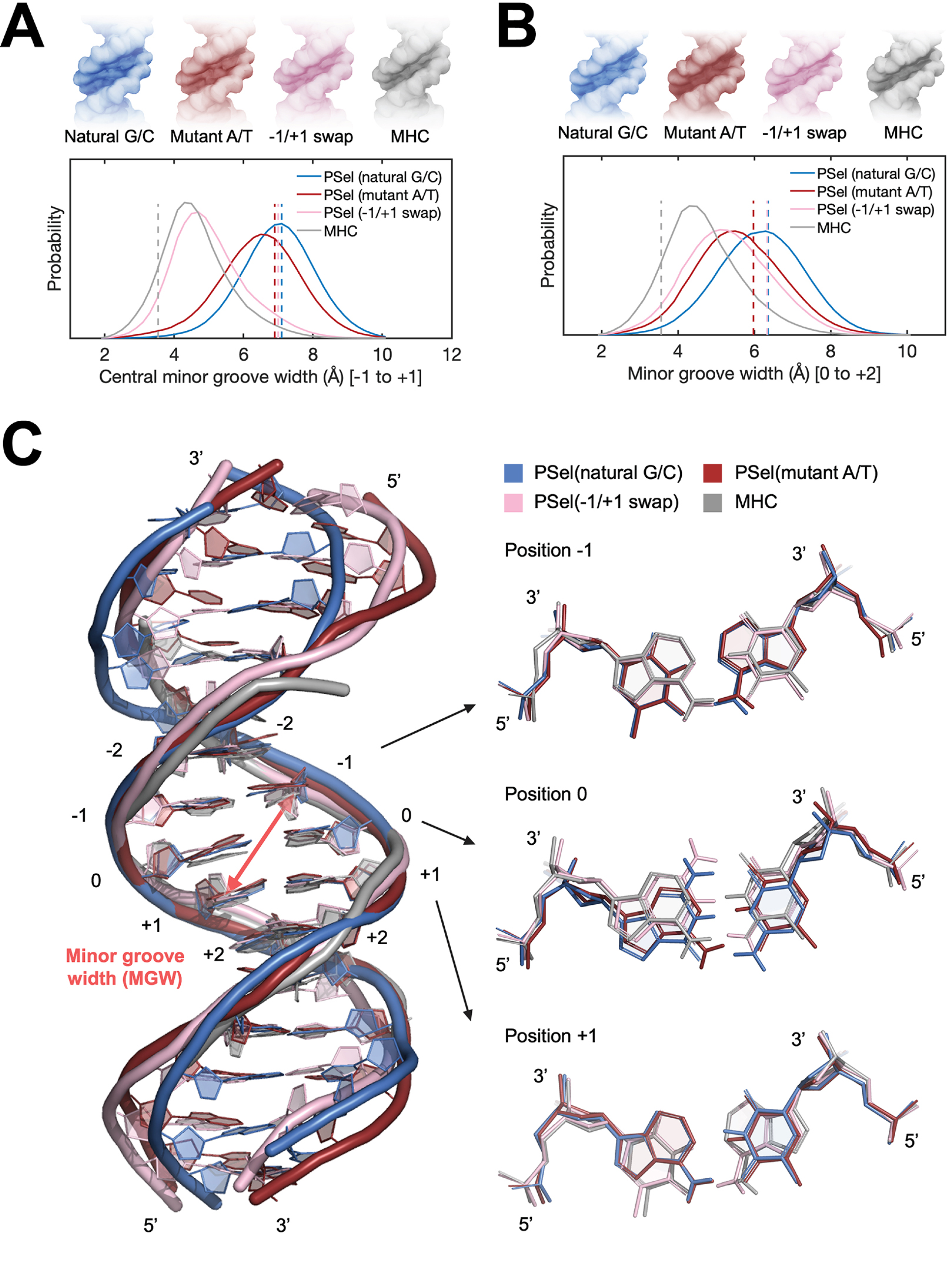
MD simulations for free κB DNAs. (A-B) Statistical MGW over the aggregated 10-µs simulations of each system at (A) the central 0 position (averaged over the five levels from −1 to +1 positions) and (B) the +1 position (averaged over the five levels from 0 to +2 positions). (Upper) DNA isosurface at 0.2 isovalue (20% occupancy); (Lower) Probability distribution of MGW. Dashed lines show the MGW in the (p52:p52)-bound crystal structures. (C) Representative structures of natural G/C-centric, mutant A/T-centric, −1/+1 swap PSel-κB DNAs and MHC-κB DNA revealed from MD simulations. (Left) Superimposed structures showing the narrowed central minor groove on −1/+1 swap DNAs; (Right) Representative conformations of bps at −1, 0 and +1 positions revealed from MD simulations. MGW was calculated with Curves+ (Blanchet et al., 2011, Lavery et al., 2009).

The simulations also reveal a narrowed minor groove of the A/T-centric DNA at +1 position compared to the corresponding G/C-centric DNA with the same flanking bps, i.e. PSel (mutant A/T-centric) compared to PSel (natural G/C-centric), and MHC compared to PSel (−1/+1-swap) (Fig. 4B). The A:T bp at 0 position shows large shear, buckle and opening, forcing a register towards the minor groove in the curvature of free A/T-centric DNAs (Supplemental Fig. S4A). This conformation orients the thymine towards the minor groove and might have caused the decrease in MGW at +1 position. Collectively, having A:T at N position or T:A at N+2 position is likely to intrinsically reduce the MGW at N+1 position. Our finding is in line with the observations of compressed minor groove of free κB-33 DNA or the bending into minor groove from continuous A:T in A-tract DNAs (Barbič et al., 2003).

Comparison of MD simulations and crystal structures suggests that upon binding to p52 the −1/+1 swap DNA experiences more disruptive conformational change than the natural G/C-centric PSel-κB (Fig. 4A; Supplemental Fig. S4B). The binding at the central part of both DNAs is symmetrically facilitated through the H-bonds between two residues (Lys143 and Tyr55) from each monomer of p52 and the phosphate of nucleotides at −1 and +1 and T-shaped π-stacking interactions between DNA bases and Tyr55. These two residues in the two p52 monomers are positioned further apart in the PSel-κB DNA complex compared to the MHC-κB DNA complex. Unlike the natural G/C-centric DNA, the bindings on both strands of −1/+1 swap DNA draw the bound thymine in the opposite direction of the minor groove, breaking the intra-bp H-bonds and severely distorts the central bps (Supplemental Fig. S4B). It appears that p52:p52 homodimer adopts a specific conformation with a nearly fixed inter-monomer distance upon the binding to κB DNAs, such that it tears the central part of −1/+1 swap DNA into a favored binding conformation. Overall, the comparative analysis of MD simulations and crystal structures suggests that the p52:p52 homodimer induces the least amount of conformational changes on κB DNA with an intrinsically widened minor groove.

### The p52 homodimer recognizes κB DNAs with different thermodynamic features

Structural analysis described above did not provide a strong correlation between the conformational states of (p52:p52)-DNA complexes and the transcriptional output. We next tested whether p52:p52 homodimer binds to the natural G/C-centric, mutant A/T-centric, and −1/+1 swap PSel-κB, as well as MHC-κB DNAs with different mechanisms and/or affinities. In all cases, long p52 protein (aa 1-398) as well as long DNA were used. Isothermal titration calorimetry (ITC) reveals that p52:p52 binds all three PSel-κB DNA variants with similar binding affinities (K_d_) of approximately 70 nM whereas it binds MHC-κB DNA more than 2-fold tighter (Fig. 5A-D). However, binding of p52 to the natural G/C-centric PSel-κB DNA is associated with a large increase in entropy (ΔS), while the binding to the MHC-κB DNA showed a large decrease in entropy. On the other hand, the binding to the MHC and mutant A/T-centric PSel DNAs showed a much larger decrease in enthalpy (ΔH). These results suggest that the binding of p52:p52 homodimer to the G/C-centric κB DNA is favored by entropy, whereas the binding to the A/T-centric DNA is driven by enthalpy.

**Figure 5.**
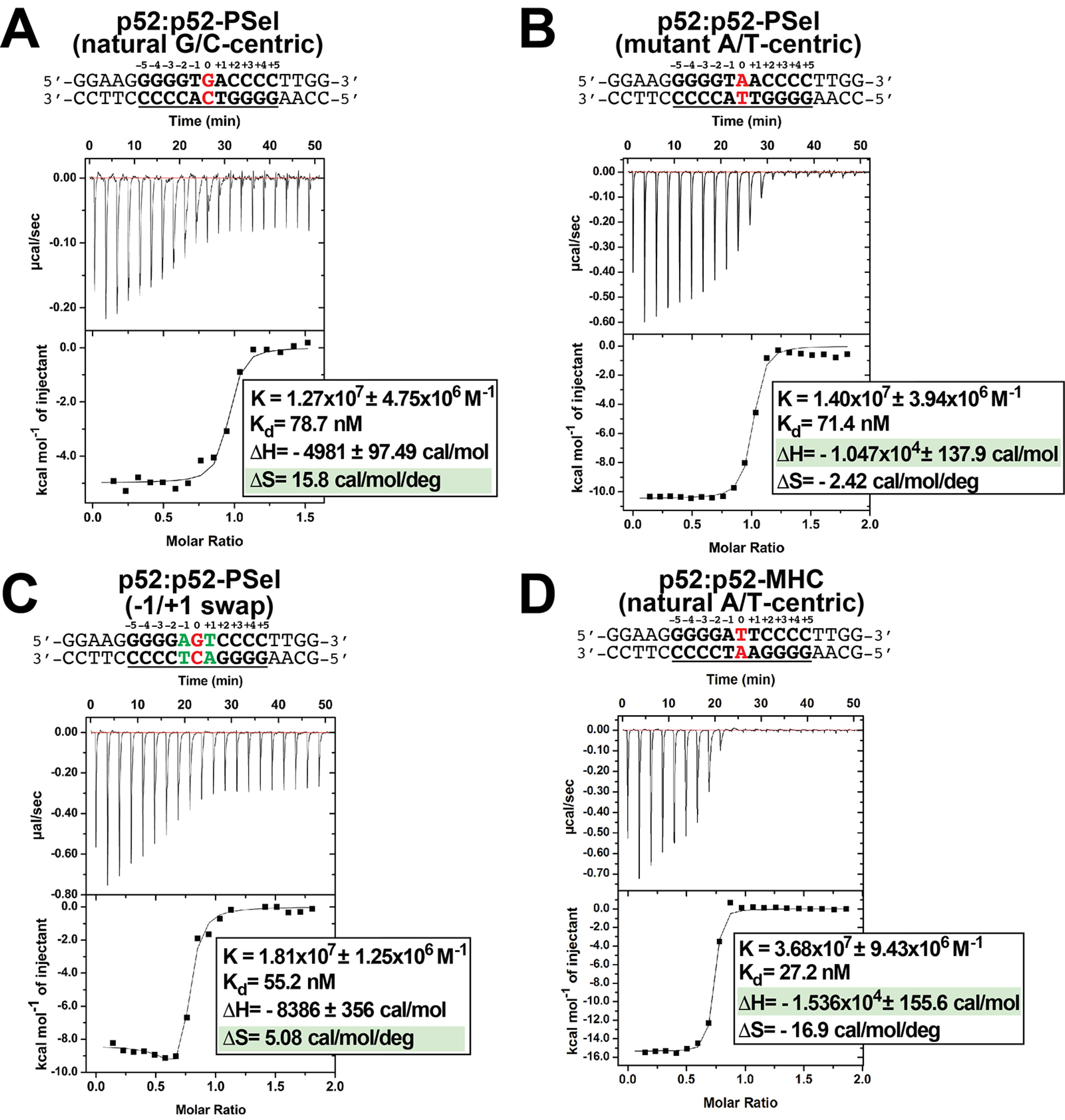
p52 binds κB DNAs with different thermodynamic features. (A-D) Calorimetric titration data showing the binding of recombinant p52:p52 (aa 1-398) homodimer with (A) natural G/C-centric, (B) mutant A/T-centric, (C) −1/+1 swap PSel-κB and (D) MHC-κB DNAs. The top panel of the ITC figures represent the binding isotherms; the bottom panel shows the integrated heat of the reaction and the line represents the best fit to the data according to a single-site binding model. The determined K_d_, changes of enthalpy and entropy are shown on the bottom panel.

To test if this mechanism is general to other κB DNAs, we also determined the thermodynamic parameters for p52 binding to Skp2-κB DNA. Skp2-κB DNA is present in the promoter of S-phase kinase-associated protein 2 (Skp2) and it is also regulated by the p52:p52 homodimer and Bcl3 (Supplemental Fig. S5A) (Barre and Perkins, 2007, Wang et al., 2012). Skp2-κB DNA is another natural G/C-centric (5’-GGGG<u>AGT</u>TCC-3’) κB DNA but with the presence of A:T and T:A bp at +1 and −1 positions, the same as the −1/+1 swap PSel DNA in the central region. Skp2 also has a very different 4bp half-site, TTCC, at the +1 to +4 positions. The K_d_ as well as relative contributions of entropy and enthalpy to the binding to Skp2 and −1/+1 swap PSel DNAs are similar (Fig. 5C; Supplemental Fig. S5B). These results suggest that DNA sequence and conformational differences lead p52 to bind DNAs through different thermodynamic binding processes. However, thermodynamic binding mechanism does not fully capture the differential transcriptional output mediated by these κB DNAs.

### The p52 homodimer binds κB DNAs with different kinetic features

We next examined the binding kinetics of p52 and κB DNAs as there is mounting evidence that, separate from binding affinity, kinetic rate constants (*k*_on_ and *k*_off_) are crucial to the physiological effects of protein-ligand interactions in a variety of cellular processes (Nakajima et al., 2001, Gross and Lodish, 2006, Gonzalez et al., 2005, Markgren et al., 2002). We utilized biolayer interferometry (BLI) to study the association and dissociation rate of p52:p52 binding to various κB DNAs. Biotinylated DNAs were immobilized on the streptavidin (SA) sensors and tested with purified p52 protein. The binding kinetics differ significantly among the DNAs. The binding of more transcriptionally active natural G/C-centric PSel showed a higher association (*k*_on_) and dissociation rate (*k*_off_) than the other two variants and MHC-κB DNAs (Fig. 6A-E). Consistently, in the case of Skp2-κB DNA, the more transcriptionally active natural G/C-centric Skp2 showed a faster kinetics than its mutant A/T-centric, especially the *k*_off_ (Supplemental Fig. S6A-C).

**Figure 6.**
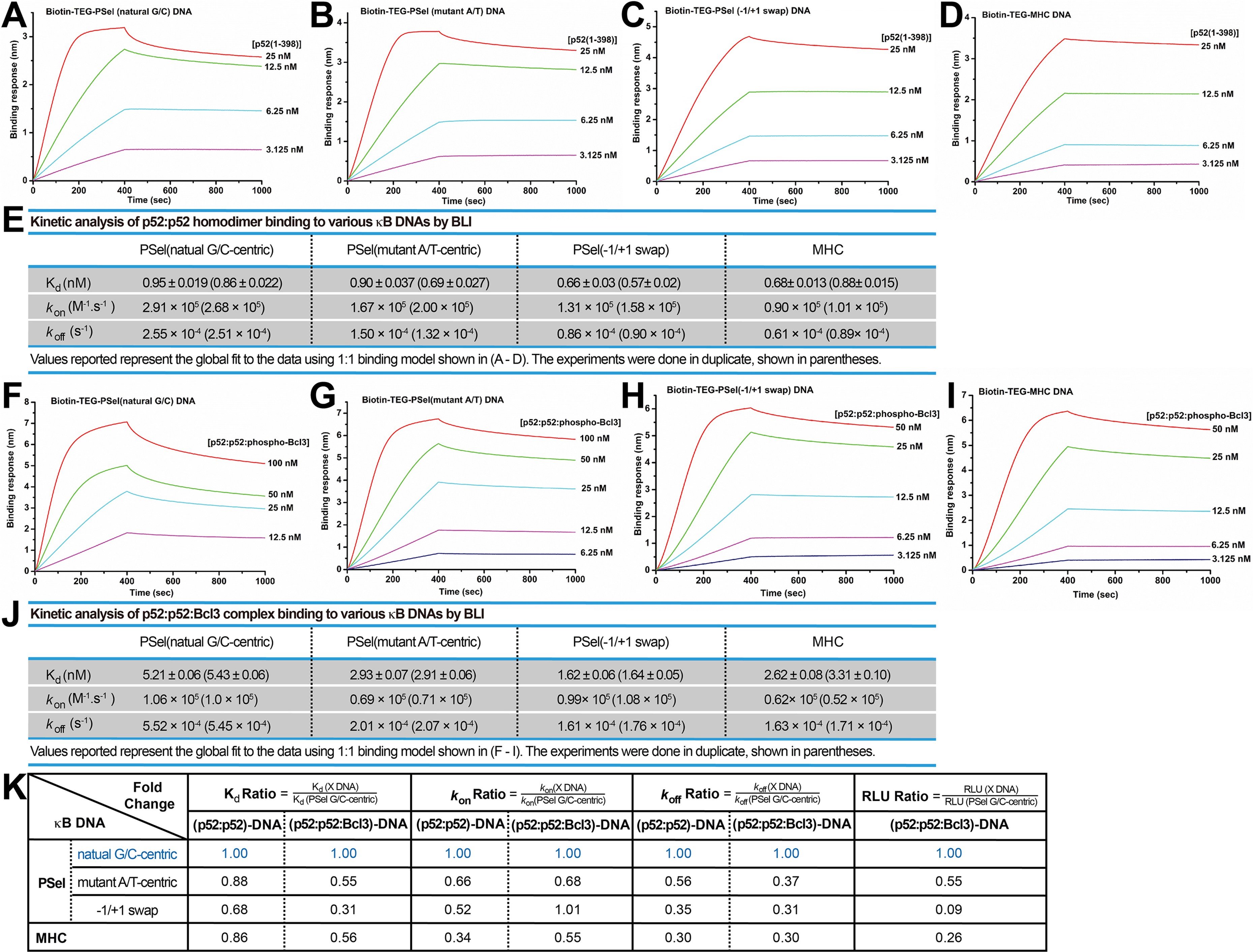
p52 binds the natural transcriptionally active G/C-centric PSel-κB DNA with faster kinetics. (A-D) Biolayer interferometry (BLI) binding analysis of p52:p52 (aa 1-398) homodimer to immobilized biotin labeled (A) natural G/C-centric, (B) mutant A/T-centric, (C) −1/+1 swap PSel-κB and (D) MHC-κB DNAs. The differences in *k*_on_ and *k*_off_ can be seen in the shapes of the association and dissociation curves. Each experiment was done in duplicate and one representative set of curves is shown. (E) Table showing the kinetic analysis in (A-D). (F-I) BLI binding analysis of p52:p52:Bcl3 complex to immobilized biotin labeled (F) natural G/C-centric, (G) mutant A/T-centric, (H) −1/+1 swap PSel-κB and (I) MHC-κB DNAs. Each experiment was done in duplicate and one representative set of curves is shown. (J) Table showing the kinetic analysis in (F-I). (K) Table summarizing the fold change of K_d_, *k*_on_ and *k*_off_ with respect to the more transcriptionally active G/C-centric PSel-κB DNA. The average values of the duplicated kinetics data in (A-J) and the relative reporter activities in RLU from Fig. 1A-B were used for ratio calculations. The numbers for the greater reporter active G/C-centric PSel are shown in blue.

We further determined the *k*_on_ and *k*_off_ of the transcriptionally competent p52:p52:Bcl3 complex binding to PSel-κB DNA variants by BLI. In agreement with our previous study, only the recombinant phospho-mimetic Bcl3 from E. coli forms ternary complex with p52:p52 homodimer and κB DNA (Wang et al., 2017). Both recombinant WT and phospho-mimetic Bcl3 protein (S33/114/446E mutant) interact with p52 with similar kinetics (Supplemental Fig. S1F, S7A-C); however, the p52:p52:WT-Bcl3 complex does not bind DNAs (Supplemental Fig. S7D-E). The binding of p52:p52:phospho-Bcl3 with the natural G/C-centric PSel DNA exhibited both higher *k*_on_ and *k*_off_ (Fig. 6F-J). Similarly, the binding with the natural G/C-centric Skp2 DNA also showed a higher *k*_off_ comparing to its A/T-centric mutant (Supplemental Fig. S6D-F).

Overall, the binding kinetics of p52:p52 homodimer alone vs. p52:p52:Bcl3 complex follows the same trend. Moreover, a comparison of binding affinity, association and dissociation rates with respect to the more transcriptionally active PSel and Skp2-κB sites shows a correlation between transcriptional output and the dissociation rate. The slower the *k*_off_, the lower the reporter activities for both (p52:p52)-DNA and (p52:p52:Bcl3)-DNA complexes (Fig. 6K; Supplemental Fig. S6G). Therefore, transcriptional activity may have a closer link to the binding kinetics rather than the thermodynamic stability of the complex.

## Discussion

### The p52 homodimer recognizes κB DNA using a mode distinct from other NF-κB dimers

Double-stranded DNA helices are not static entities that simply present themselves to proteins and assemble into multiprotein complexes at specific sequences. The DNA duplex is intrinsically dynamic on many levels and time scales in cells. The movement of DNA through its different conformational states is continuous and is influenced by, but not completely dependent upon, its nucleotide sequence. Structures presented in this study show that the conformations of all three PSel-κB DNA variants bound to the long p52:p52 homodimer are similar but are distinct from all previously known complexes between κB DNAs and six other NF-κB dimers (p50:p50, p50:RelA, p50:RelB, RelA:RelA, c-Rel:c-Rel and p52:RelB). It was noted earlier a compressed minor groove in the central region of the DNA is a key feature of NF-κB-DNA complexes. The minor groove at the central three positions is significantly widened in all three complexes presented here. However, MD simulations show free DNAs exist in distinct preferred conformations, which appears to be adjusted by p52 into a unique shape for recognition. And notably, the more transcriptionally active natural PSel-κB DNA appears to maintain similar conformational and dynamic states in free and bound forms.

The current structures also demonstrate a correlation between the p52 protein length and the conformation of the κB DNA. The GRR region of the p52 protein, which was not included in any previous NF-κB structural studies, seems to play an important role. However, no electron density was observed for the GRR region in the current structures. Future studies are needed to fully understand the role of the GRR region in (p52:p52)-DNA complex conformation and the interaction with cofactor Bcl3.

### Binding affinity does not fully capture the transcriptional activity

To determine if binding affinity is related to transcriptional activity, we measured the affinity of all complexes under equilibrium condition. Surprisingly, but consistent with our previous report (Mulero et al., 2018), we found no correlation between affinity and transcriptional activity. The p52:p52 homodimer binds to MHC-κB with the highest affinity but it is not a transcriptional activation competent complex. Interestingly, our analysis reveals that p52:p52 homodimer uses different paths to bind κB DNAs ranging from purely entropic for the natural PSel, to exclusively enthalpic for MHC, and to mixed entropic-enthalpic for mutant A/T-centric and −1/+1 swap PSel DNAs. The entropy-driven processes are linked to faster binding kinetics: p52:p52 homodimer binds to natural G/C-centric PSel DNA with both faster association and dissociation rates. The most populated conformation of the free G/C-centric PSel DNA as revealed by MD simulation is similar to the one observed in the crystal structure of the complex suggesting this DNA’s conformation does not undergo significant changes upon protein binding. Thus, in the complex between p52:p52 and natural G/C-centric DNA, both the DNA and protein most likely preserve their native states. This could account for the positive entropy and faster *k*_on_. However, possibly protein-DNA contacts in such a complex are sub-optimal resulting in their faster dissociation. In contrast, p52:p52 complexes with the mutant A/T-centric or −1/+1 swapped DNAs likely involve rigidification of protein-DNA contacts, requiring some structural reorganizations in both molecules, results in more enthalpically stable complexes and slower association and dissociation rates. Combination of MD simulations and structural studies supports this model as PSel (mutant A/T) and (−1/+1 swap) DNAs undergo conformational changes from free to bound states. Our observations are consistent with other studies which showed rapid association and dissociation is favored by entropy, whereas slow association and dissociation is guided by enthalpy (Baerga-Ortiz et al., 2004).

### Ideas and Speculation: Transcriptional regulation via kinetic discrimination

Understanding transcriptional regulation has attracted many researchers since the discovery of the *lac* operon. One of the most intriguing questions scientists are working to resolve is the mechanism of transcriptional regulation by the specific DNA response elements. Affinity regulation by different target DNA sequences for a TF has long been thought to be the dominant mode of regulation imposed by such DNA sequences. Indeed, in many cases of eukaryotic transcription differential affinity has been shown to be critical (Sekiya et al., 2009). TFs are also known to bind free DNA or nucleosome with distinct kinetics (Donovan et al., 2019). But none of these studies has established a direct correlation between binding kinetics and transcriptional regulation in eukaryotes. Many biological systems have been studied in detail with the roles of binding kinetics in regulation evaluated. For instance, receptor ligand interactions, T cell activation and potency of bacterial toxins are guided by the half-life of key complexes (Corzo, 2006, Gonzalez et al., 2005, Gross and Lodish, 2006, Nakajima et al., 2001).

Of all six κB DNAs tested, the natural G/C-centric PSel and Skp2 DNAs showed greater transcriptional activation. We found that a slower dissociation rate or longer residence time is linked to reduced transcriptional activation. Although the relationship between the dissociation rates and transcription activities is shown for only six tested binding sites, this relationship is preserved for (p52:p52)-DNA complexes and to a lesser extent for (p52:p52:Bcl3)-DNA complexes.

The rate constants obtained in our *in vitro* assays probably are not the same *in vivo*, where many other factors will have an impact on binding kinetics. However, the relative rates clearly suggest that the persistent presence of p52 on DNA gives rise to less transcription. Work presented here hints at a link between the DNA binding kinetics of a TF and its interaction with coactivators and corepressors. We previously showed that the p52:p52:Bcl3 complex preferentially recruits HDAC3 when it remains bound to an A/T-centric κB site (Wang et al., 2012), it is possible that the slower dissociation rate or longer residence time of p52:p52:Bcl3 on the A/T-centric κB site described in this study matches the slower on rate of the corepressor to p52:p52:Bcl3 bound to A/T DNA. That is, the A/T-centric κB DNA:p52:p52:Bcl3 complex remains stable for long enough to give the HDAC3 corepressor complex enough time to stably interact with it. In addition, the binding of other TFs to the promoters/enhancers of target genes inevitably impacts on the coactivator/corepressor regulation by the (p52:p52)-DNA complexes. Future experiments aimed at coactivator and corepressor interaction rates within the context of chromatinized DNA are needed to verify the validity of the kinetic model for DNA element sequence-specific gene regulation.

In summary, our studies have revealed a novel conformation for κB DNA in complex with NF-κB and a new organization of an NF-κB dimer. More importantly, our work provides a new insight into the mechanism of differential thermodynamics and kinetics of NF-κB-DNA binding. DNA response elements with only one or two bp variations could provoke drastically different kinetic and thermodynamic effects. Future experiments will help us fully understand how such binding processes result in transcription activation or repression.

## Materials and Methods

### Protein expression and purification

Recombinant non-tagged human p52 (1-398) and (1-327) was expressed and purified from *Escherichia coli* Rosetta (DE3) cells. Rosetta (DE3) cells transformed with pET-11a-p52 (1-398) or (1-327) were cultured in 2 L of LB medium containing 50 mg/mL ampicillin and 34 mg/mL chloramphenicol at 37 °C. Expression was induced with 0.2mM Isopropyl β-D-1-thiogalactopyranoside (IPTG) at OD_600_ 0.5-0.6 for 3 hours. Cells were harvested by centrifugation, suspended in 40 mM Tris-HCl (pH 7.5), 100 mM NaCl, 10 mM β-ME, 1 mM PMSF, and lysed by sonication. Cell debris was removed by centrifugation (20,000 g for 30 min). Clarified supernatant was loaded onto Q-Sepharose FF column (GE Healthcare). Flow-through fraction was applied to SP HP column (GE Healthcare). The column was washed with 40 mM Tris-HCl (pH 7.5), 200 mM NaCl; 10 mM β-ME, and the protein was eluted by the same buffer containing 400 mM NaCl. p52 was concentrated and loaded onto the gel filtration column (HiLoad 16/600 Superdex 200 pg, GE Healthcare) pre-equilibrated with 10 mM Tris-HCl (pH 7.5), 100 mM NaCl; 5 mM β-ME. Peak fractions were concentrated to ∼10 mg/mL, flash frozen in liquid nitrogen and stored at −80°C. His-Bcl3 (1-446) WT and phospho-mimetic mutant was expressed in *Escherichia coli* Rosetta (DE3) cells by induction with 0.2 mM IPTG at OD_600_ 0.4 for 8 hours at 24°C. Cell pellets of 2 L-culture of Bcl3 alone or together with 1 L-culture of p52 (for p52:Bcl3 complex) were resuspended together in buffer containing 20 mM Tris-HCl (pH 8.0), 300 mM NaCl, 25 mM imidazole, 10% glycerol, 10 mM β-ME, 0.1 mM PMSF and 50μL protease inhibitor cocktail (Sigma) and then purified by Ni Sepharose (HisTrap HP, GE) followed by anion exchange column (Q Sepharose fast flow, GE). The protein complex further went through HiTrap Desalting Column (GE) to exchange buffer before BLI assays.

### Crystallization, data collection, and structure determination

Annealed DNA duplex was mixed in 20% molar excess with the pure protein.

The crystals of the p52(aa 1-398):PSel(G/C-centric) 18bp complex were obtained by the sitting-drop vapor diffusion method at 20 °C with a reservoir solution containing 0.1 M sodium malonate (pH 4.0), 0.2 M CsCl and 5% (w/v) PEG 3350.

The crystals of the p52(aa 1-398):PSel(A/T-centric) 18bp complex, and the p52(aa 1-398):PSel(−1/+1 swap) 18bp complex were obtained by the sitting-drop vapor diffusion method at 20 °C with a reservoir solution containing 0.1 M sodium malonate (pH 4.0), 50 mM CsCl and 2.5% (w/v) PEG 3350.

The crystals of the p52(aa 1-327):Psel(A/T-centric) 13bp complex were obtained by the sitting-drop vapor diffusion method at 20 °C with a reservoir solution containing 50 mM MES (pH 6.0), 10 mM MgCl_2_ and 10% (w/v) PEG 3350.

Before data collection, all crystals were briefly soaked in their original crystallization solution with 20% (v/v) ethylene glycol. All crystals were flash frozen in liquid nitrogen for diffraction screening and data collection at 100 K. X-ray diffraction data were collected at beamline BL19U1 at Shanghai Synchrotron Radiation Facility. The initial solution was obtained by molecular replacement using Phaser (McCoy et al., 2007) with p52-MHC DNA complex (Cramer et al., 1997) as the search model. The structure was further refined through an iterative combination of refinement with Refmac5 (Murshudov et al., 2011) and manual building in the Coot program (Emsley and Cowtan, 2004, Emsley et al., 2010). The crystallographic information is summarized in Table 1.

### MD simulation

All MD simulations were carried out in GROMACS 2020.6 (Lindahl et al., 2020) with Amber14sb force field (Maier et al., 2015) and OL15 parameters for DNA (Zgarbová et al., 2015). Crystal structures of κB/κB-like DNAs were extracted from the resolved p52-bound structures, where the MHC DNA was retrieved from RCSB PDB database (PDB 1A3Q) (Cramer et al., 1997). In each system, κB DNA was placed in the center of a dodecahedron box with a 12-Å margin, solvated with TIP3P water (Jorgensen et al., 1983), and ionized with 0.1 M NaCl. Energy minimization was performed until the maximum force of system was below 1,000 kJ·mol^-1^·nm^-1^. The minimized system was then equilibrated in a NVT ensemble for two 1-ns stages, positionally restraining the DNA heavy atoms with a force constant of 20,000 kJ·mol^-1^·nm^-2^ and 10,000 kJ·mol^-1^·nm^-2^, respectively. Subsequently, the system was subjected to a 6-ns position-restrained NPT equilibration, with the force constant gradually reduced from 10,000 kJ·mol^-1^·nm^-2^ to 400 kJ·mol^-1^·nm^-2^. Finally, five replicas of 2-µs unrestrained production simulations were run for each well-equilibrated DNA system, resulting in an aggregated 10-µs trajectory for each system. In all simulations, van der Waals forces were smoothly switched to zero from 9 Å to 10 Å. Electrostatics were calculated using the particle mesh Ewald (PME) method (Darden et al., 1993) with a cutoff of 10 Å. A velocity-rescaling thermostat (Bussi et al., 2007) was employed for the temperature coupling at 300 K, whereas pressure coupling at 1 bar was implemented by a Berendsen barostat (Berendsen et al., 1984). All bonds involving H atoms were constrained using the LINCS algorithm (Hess et al., 1997).

Occupancy of DNA was calculated over the aggregated integrated 10-µs trajectories using the VolMap tool in VMD (Humphrey et al., 1996). The clustering analyses were conducted within GROMACS packages using GROMOS method. Representative structures of DNA and bp at position −1, 0, +1 were obtained from the centroid structures of top clusters and rendered with PyMOL (Schrodinger, 2015). The hydrogen bonds were calculated using PyMOL with default standard (heavy atom distance cutoff of 3.6 Å and angle cutoff of 63°). The bp and groove parameters were measured via Curves+ (Lavery et al., 2009, Blanchet et al., 2011), with the uncertainty represented by the standard error of the mean (SEM) computed from the five replica simulations of a given system.

### Isothermal titration calorimetry (ITC) assays

ITC measurements were carried out on a MicroCal iTC200 (Malvern Inc.) at 25°C. The ITC protein sample p52 (1-398) went through desalting column (HiTrap desalting, GE Healthcare) to freshly made ITC buffer containing 20 mM Tris-HCl (pH 8.0), 100 mM NaCl, 1mM dithiothreitol (DTT). 35 µM p52 (1-398) protein (in cell) was titrated with 300 µM DNAs (in syringe). A time interval of 150 seconds between injections was used to ensure that the titration peak returned to the baseline. The titration data were analyzed using the program Origin7.0 and fitted by the One Set of Site model.

### Biolayer interferometry (BLI) assays

The kinetic assays were performed on Octet K2 (ForteBio) instrument at 20°C with shaking at 1000 RPM. The streptavidin (SA) biosensors were used for protein-DNA interactions and were hydrated in BLI buffer containing 20 mM Tris-HCl (pH 8.0), 100 mM NaCl, 1 mM DTT and 0.02% (v/v) Tween-20. All DNAs used were 20-mer in length and biotin-triethyleneglycol (TEG) labelled. The DNAs were loaded at 50 nM for 300 sec prior to baseline equilibration for 60 sec in the BLI buffer. Association of p52:p52 (aa 1-398) or p52:p52:Bcl3 complex in BLI buffer at various concentrations was carried out for 400 sec prior to dissociation for 600 sec. The Ni-NTA biosensor were used for protein-protein interactions and were hydrated in BLI buffer containing 20 mM Tris-HCl (pH 8.0), 200 mM NaCl, 5% glycerol, 1 mM DTT and 0.02% (v/v) Tween-20. His-tagged-Bcl3 was loaded at 500 μg/mL for 90 sec prior to baseline equilibration for 180 sec in the BLI buffer. Association of p52 in BLI buffer at various concentrations was carried out for 240 sec prior to dissociation for 360 sec. All data were baseline subtracted and analyzed in ForteBio data analysis software using a global fitting to a 1:1 binding model. The experiments were done in duplicate.

### Luciferase Reporter Assays

HeLa cells were transiently transfected with Flag-p52(1-415) together with Flag-Bcl3(1-446) expression vectors or empty Flag-vector, and the luciferase reporter DNA with specific κB DNA promoter (Wang et al., 2012). The total amount of plasmid DNA was kept constant for all assays. Transient transfections were carried out using Lipofectamine 2000 (Invitrogen). Cells were collected 48 hours after transfection. Luciferase activity assays were performed using Dual-Luciferase Reporter Assay System (Promega) following the manufacturer’s protocol. Data are represented as mean standard deviations (SD) of three independent experiments in triplicates.

## Data Availability

The atomic coordinates have been deposited in the Protein Data Bank, www.wwpdb.org (PDB ID codes 7CLI, 7VUQ, 7VUP and 7W7L).

## Acknowledgments

We thank the Proteomics, Metabolomics and Drug Development (PMDD) core at Faculty of Health Sciences for providing the ITC machine for the thermodynamic assays. We thank the staffs from BL19U1 beamline of National Facility for Protein Science in Shanghai (NFPS) at Shanghai Synchrotron Radiation Facility, for assistance during data collection. We thank Prof. Liang Tong at Columbia University for critical discussion of the manuscript. This work was supported by the Science and Technology Development Fund, Macao S.A.R. (FDCT) [project 0104/2019/A2 to V.Y.-F.W.]; the Multi-Year Research Grant from University of Macau [MYRG2018-00093-FHS to V.Y.-F.W.]. T.L. and Y.W. were supported by direct grants from the Chinese University of Hong Kong. G.G. were supported by the National Institutes of Health (NIH) [GM085490 and CA142642 to G.G.].

## Author Contributions

V.Y.-F.W. designed the experiments and supervised the project. V.M. performed the complex crystallization, structure determination and refinement. W.P. carried out all the biochemistry, thermodynamic and binding kinetic studies. T.L. performed MD simulation. V.M., W.P., T.L., Y.W., G.G. and V.Y.-F.W. analyzed the data. V.Y.-F.W., G.G., W.Y. and T.L. wrote the paper.

---

**Table 1. Summary of crystallographic information.**

## Supplemental Information

**Figure S1.**
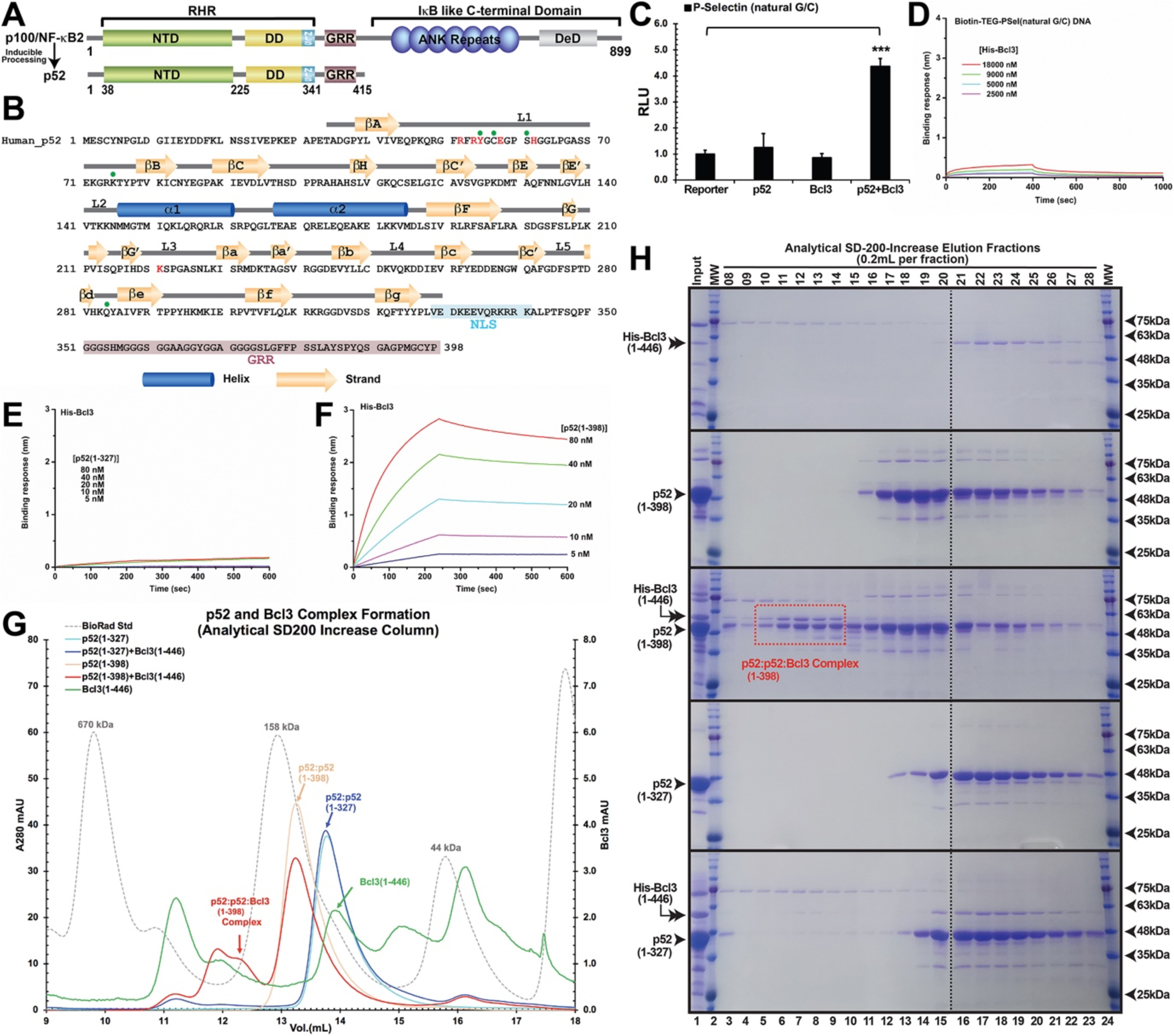
p52 and DNA crystallization. (A) Domain organization of the precursor protein p100/NF-κB2 and the processed p52. (B) Primary sequence of human p52 RHR. The secondary structures are mapped on top of the sequence. DNA base-specific contacting residues are denoted by red and backbone contacting residues are marked by green circle. NLS and GRR regions are highlighted in cyan and purple, respectively. (C) Co-expression of p52 and Bcl3, but not either one alone, activates the natural PSel luciferase reporter. The data were analyzed from three independent experiments performed in triplicate. RLU, relative luciferase unit. ***p<0.001 (t test). Error bars represent SD. (D) Biolayer interferometry (BLI) binding analysis of His-tagged-Bcl3 protein to immobilized biotin labeled natural PSel-κB DNA; the result indicated Bcl3 does not interact with κB DNA without p52 protein. (E-F) BLI binding analysis of (E) short p52:p52 (aa 1-327) and (F) long p52:p52 (aa 1-398) protein to immobilized His-tagged-Bcl3. The results showed only the long p52:p52 (aa 1-398) interacts with Bcl3 in (F). (G) Analytical Superdex-200-Increase size exclusion chromatography elution profile showing the long p52:p52 (aa 1-398) but not the short p52:p52 (aa 1-327) homodimer forms complex with Bcl3. (H) SDS-PAGE analysis indicating the p52:p52:Bcl3 complex formation in (G).

**Figure S2.**
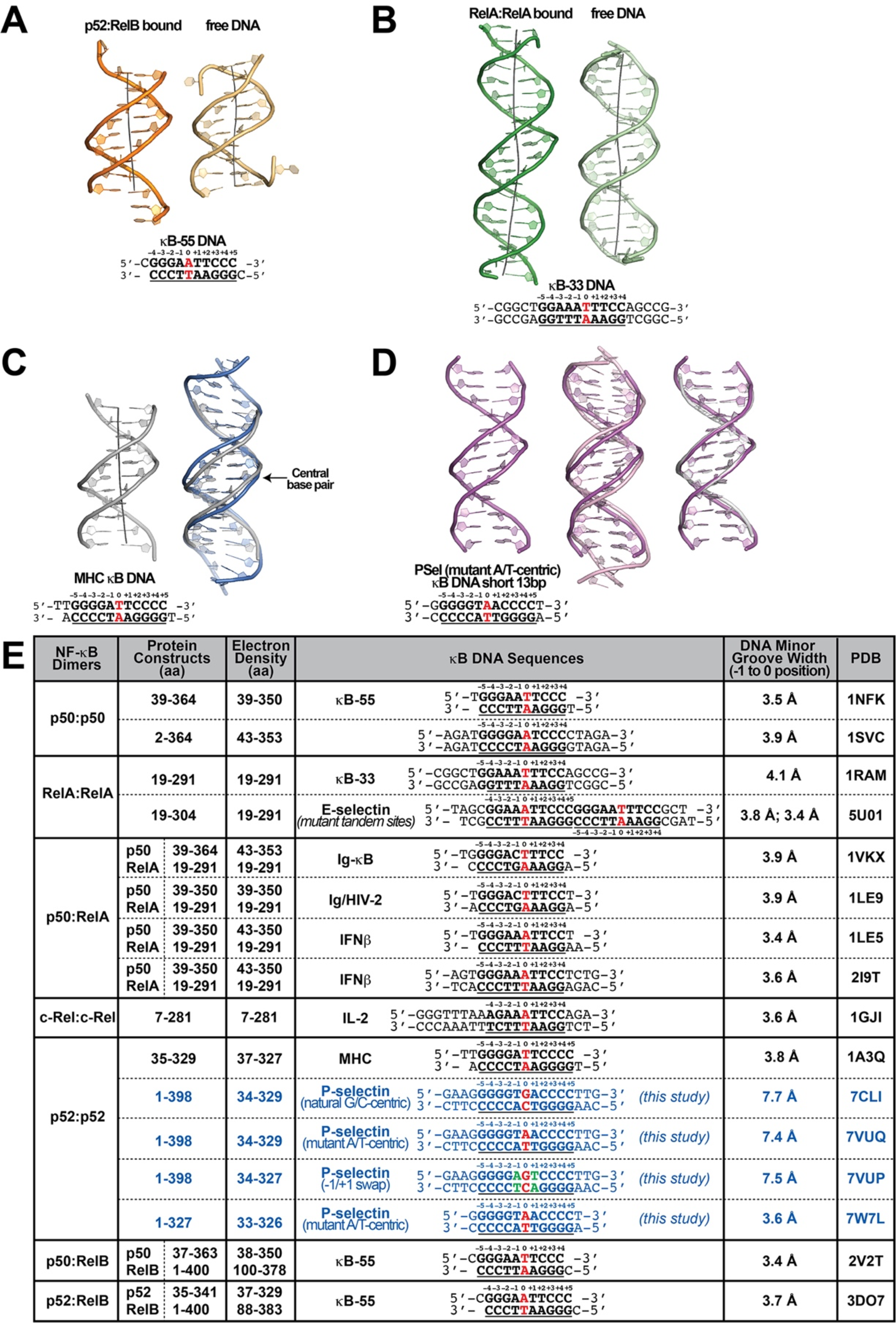
κB DNA conformations. (A) Structure of the κB-55 DNA in the (Left) p52:RelB-bound and (Right) free forms; free κB-55 DNA structure was obtained serendipitously where two κB-55 DNA molecules were found in the crystal, with one bound to the p52:RelB heterodimer and the other remained free (Fusco et al., 2009). (B) Structure of the 20bp κB-33 DNA in the (Left) RelA:RelA-bound (Chen et al., 1998) and (Right) free forms (Huang et al., 2005). (C) Structure of (Left) the 13bp MHC-κB DNA (Cramer et al., 1997) and (Right) overlay of natural G/C-centric PSel-κB DNA (in blue) with MHC-κB DNA (in gray), showing the widened minor groove in PSel-κB DNA. (D) Structure of (Left) short 13bp PSel (mutant A/T) κB DNA duplex in the co-crystal structure with short p52:p52 (aa 1-327) (in pink); overlay of 13bp PSel (mutant A/T) DNAs (in pink) with (Middle) the long 18bp PSel (mutant A/T) (in light pink), and (Right) the 13bp MHC-κB DNA (in gray). The DNA bps as observed in the co-crystal structures are shown in filled sticks. The view is onto the central minor groove. The nucleotide sequences used in co-crystallization are shown at the bottom, with κB DNA underlined and numbering scheme indicated above.(E) Table showing various nucleotide sequences used in co-crystallization with different NF-κB dimers The various NF-κB protein constructs used are also indicated. The κB DNAs are shown in bold, bases making contacts with NF-κB proteins are underlined, and the numbering scheme are indicated. The central bps are highlighted in red. The p52 protein construct and DNA sequences used in the current study are in blue. The MGW(s) from position −1 to 0 are listed. Geometrical parameters were calculated with Curves*+* (Blanchet et al., 2011, Lavery et al., 2009).

**Figure S3.**
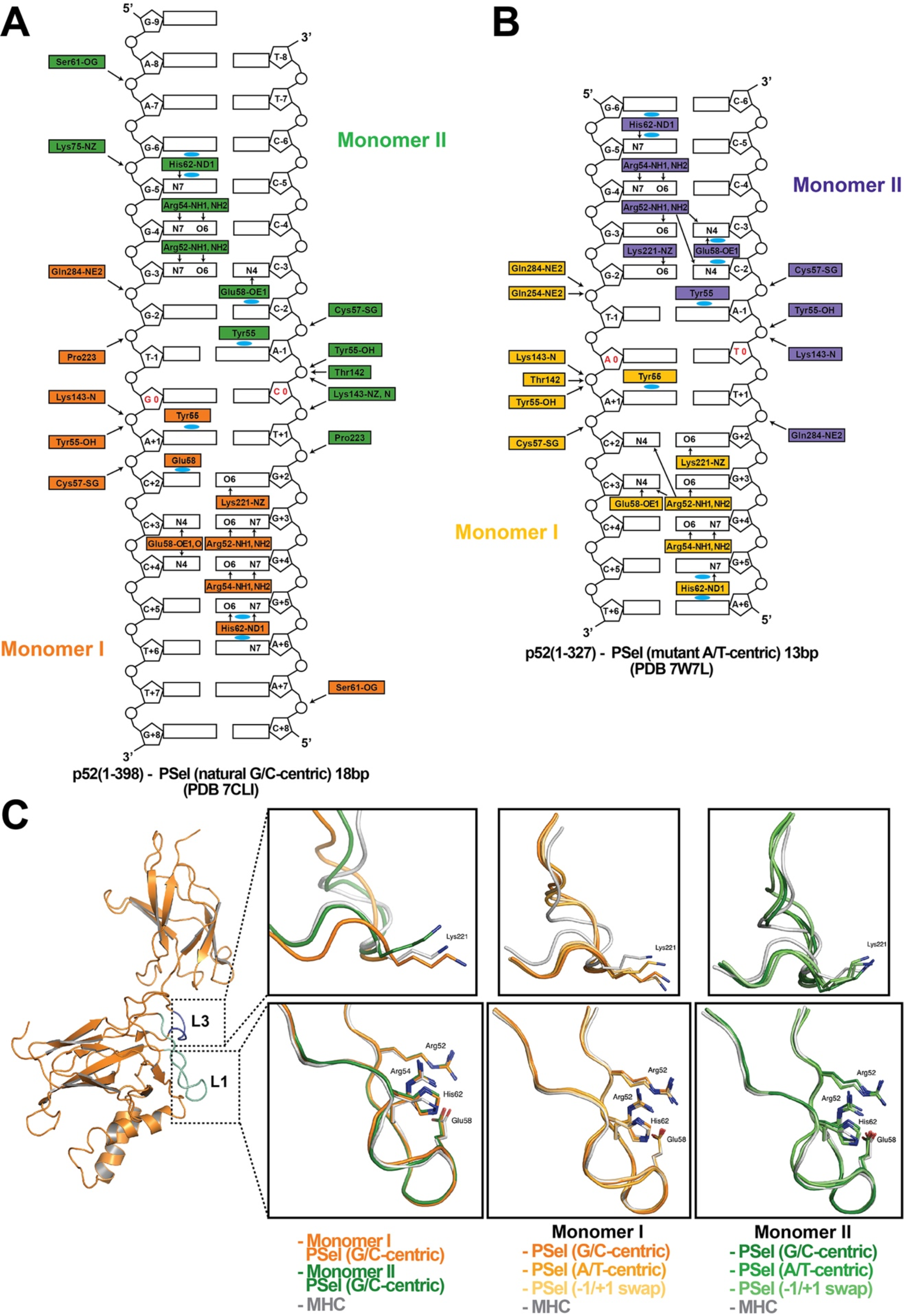
Asymmetric p52 monomers. (A-B) Schematic representation of the DNA contacts made by (A) the long p52:p52 (aa 1-398) with the 18bp natural PSel-κB DNA (PDB 7CLI); and (B) the short p52:p52 (aa 1-327) with the 13bp PSel(mutant A/T)-κB DNA (PDB 7W7L). Two colors indicate two different monomers within the complex. Arrows indicate H-bonds; cyan circles indicate van der Waals interactions. Arg52(s) from both monomers make cross-strand DNA contacts in (B) but not in (A). (C) Ribbon diagram of p52 monomer I, loops L1 (cyan) and L3 (blue) are highlighted. Residues in these two loops make base-specific contacts with DNA. The zoom-in views show overlays of loops L1 and L3 in (p52:p52)-PSel and (p52:p52)-MHC complexes with indicated colors.

**Figure S4.**
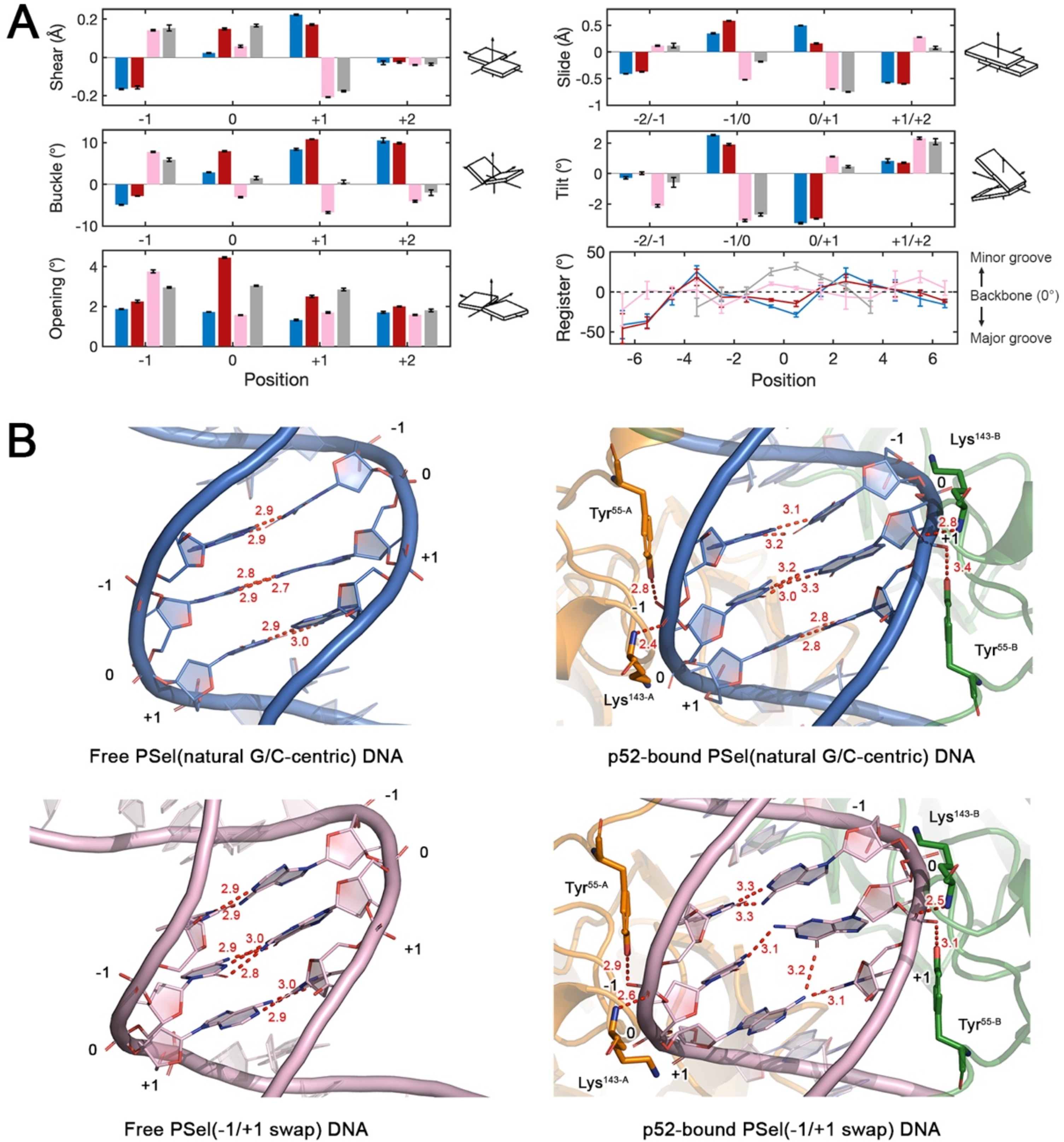
Free DNA simulations. (A) Geometric parameters at −1 to +2 positions revealed in the MD simulations. The swap of T and A at ±1 positions causes an opposite shear and buckle of the nucleotide, which leads to a slide and tilt of the central bps towards the minor groove in −1/+1 swap DNA and therefore narrows the central minor grooves; A:T has a larger shear and opening at the 0 position compared to G:C which might have led to the decrease in minor groove width at +1 position. Geometrical parameters and the helical axes were calculated with Curves+ (Blanchet et al., 2011, Lavery et al., 2009). Corresponding schematic images of each parameter are viewed from the minor groove and shown in the positive sense. Error bars represent standard error of the mean (SEM) computed from the five replica simulations of a given system.(B) Structure of natural G/C-centric and −1/+1 swap PSel-κB DNA in (Left) free forms from MD simulations and (Right) (p52:p52)-bound forms from crystal structures. Red dashed lines represent the intermolecular H-bonds formed at DNA’s central part.

**Figure S5.**
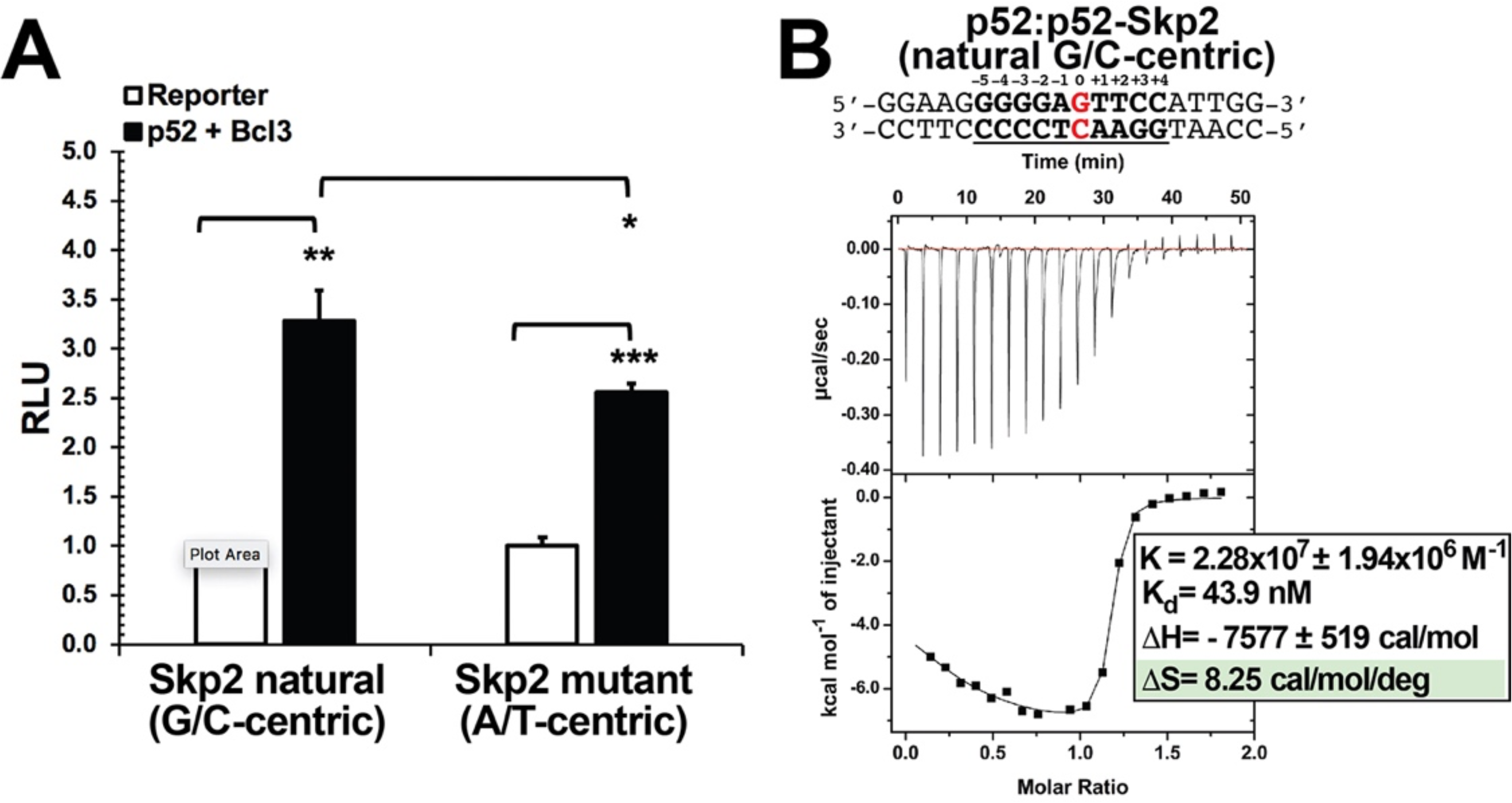
p52 interacts with Skp2-κB DNA. (A) The natural G/C-centric Skp2 luciferase reporter activity driven by co-expression of p52 and Bcl3; the corresponding A/T-centric mutant showed less transcription activity. The data were analyzed from three independent experiments performed in triplicate. RLU, relative luciferase unit. *p<0.05; **p<0.01; ***p<0.001 (t test). Error bars represent SD. (B) Calorimetric titration data showing the binding of recombinant p52:p52 homodimer with Skp2 G/C-centric κB DNA. The top panel of the ITC figures represent the binding isotherms; the bottom panel shows the integrated heat of the reaction and the line represents the best fit to the data according to a single-site binding model. The determined K_d_, changes of enthalpy and entropy are shown on the bottom panel.

**Figure S6.**
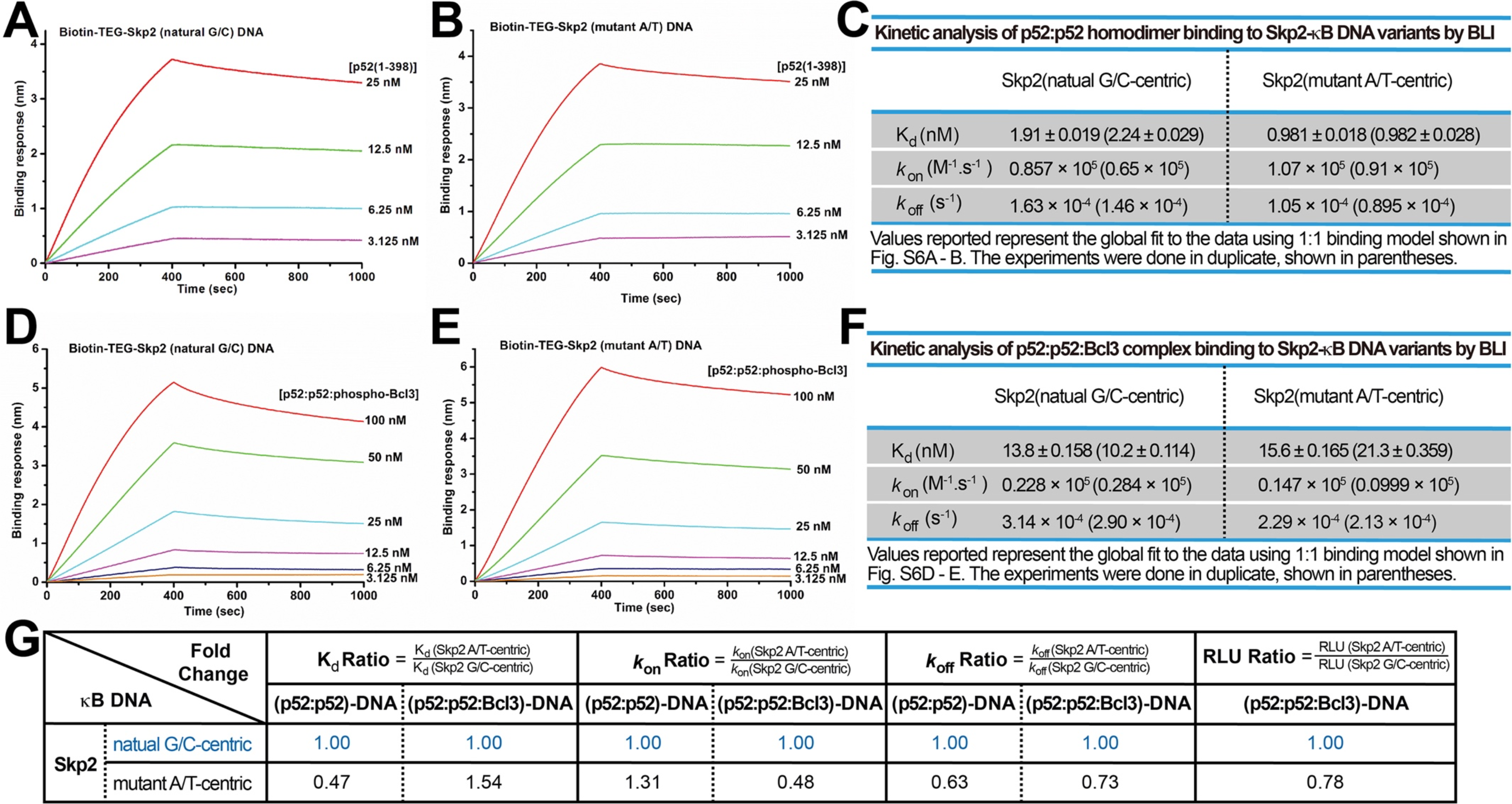
p52 and Skp2-κB DNA binding kinetics. (A-B) BLI binding analysis of p52:p52 (aa 1-398) homodimer to immobilized biotin labeled (A) Skp2 natural G/C-centric and (B) mutant A/T-centric DNAs. Each experiment was done in duplicate and one representative set of curves is shown. (C) Table showing the kinetic analysis in (A) and (B). (D-E) BLI binding analysis of p52:p52:Bcl3 complex to immobilized biotin labeled (D) Skp2 natural G/C-centric and (E) mutant A/T-centric DNAs. Each experiment was done in duplicate and one representative set of curves is shown. (F) Table showing the kinetic analysis in (D) and (E). (G) Table summarizing the fold change of K_d_, *k*_on_ and *k*_off_ with respect to the more transcriptionally active G/C-centric Skp2-κB DNA. The average values of the duplicated kinetics data in (A-F) and the relative reporter activities in RLU from Fig. S5A were used for ratio calculations. The numbers for the greater reporter active G/C-centric Skp2 DNAs are shown in blue.

**Figure S7.**
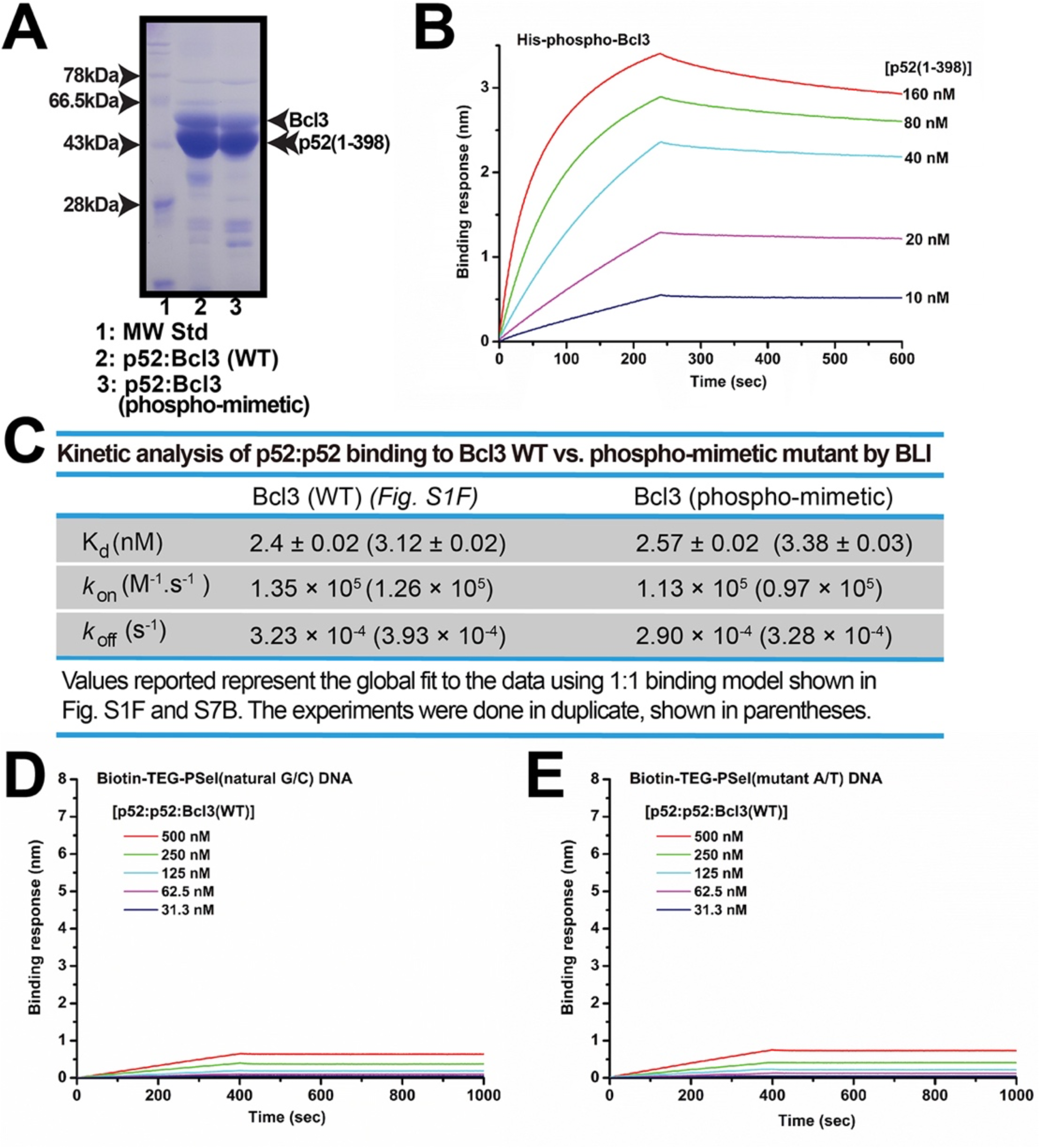
Recombinant phospho-mimetic Bcl3 forms a ternary complex with p52 and κB DNA. (A) SDS-PAGE analysis showing the p52:p52:Bcl3 WT vs. phospho-mimetic mutant complexes with similar purity. (B) BLI analysis of p52 to immobilized His-tagged-Bcl3 phospho-mimetic mutant protein. (C) Table showing the kinetic analysis in Figures S1F and S7B, suggesting p52:p52 homodimer interacts with WT and phospho-mimetic Bcl3 with similar affinity and kinetics. (D-E) BLI analysis of p52:p52:Bcl3 (WT) complex to immobilized PSel (D) natural G/C-centric and (E) mutant A/T-centric DNAs. The WT complex does not interact with either DNAs. Each experiment was done in duplicate and one representative set of curves is shown.

**Table S1.**
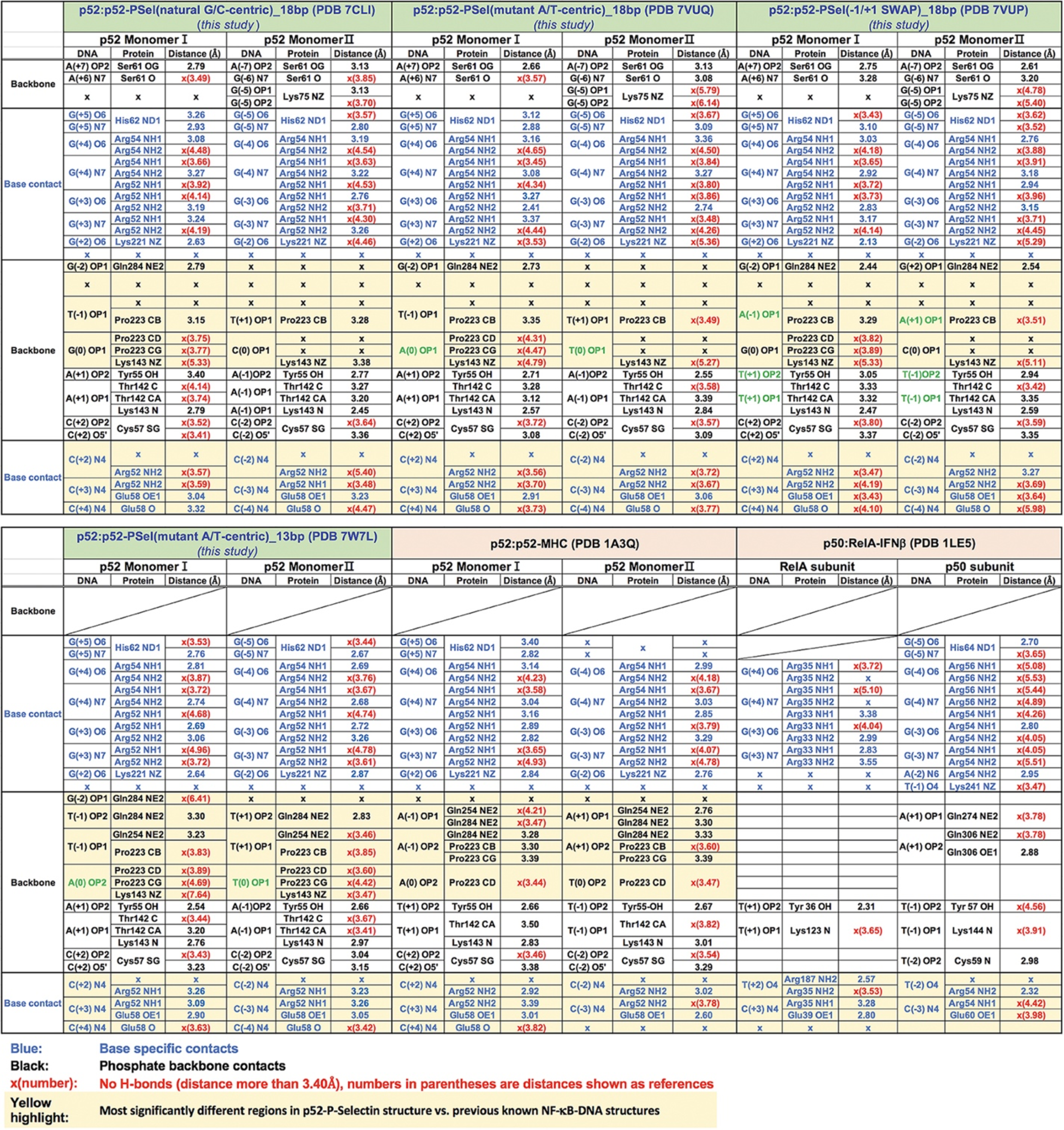
Summary of protein-DNA contacts.

